# The circadian clock gates lateral root development

**DOI:** 10.64898/2026.01.14.699582

**Authors:** Sota Nomoto, Allen Mamerto, Shiho Ueno, Akari E Maeda, Saori Kimura, Kosuke Mase, Ayano Kato, Takamasa Suzuki, Soichi Inagaki, Satomi Sakaoka, Norihito Nakamichi, Todd P. Michael, Hironaka Tsukagoshi

## Abstract

Lateral root (LR) formation remodels root architecture, yet how temporal information integrates with hormonal cues remains unclear. We show that the circadian clock component EARLY FLOWERING 3 (ELF3) acts as a temporal gatekeeper of root organogenesis in *Arabidopsis thaliana*. Time-resolved transcriptomics, imaging, and genetics reveal that ELF3 restrains hormone-induced pericycle proliferation by maintaining rhythmic expression of key regulators under callus-inducing conditions. Loss of ELF3 disrupts rhythmic gene expression, enhancing callus growth and accelerating LR development. ELF3 functions through its target LNK1 to regulate the MADS-box genes *AGL14* and *AGL20*. Notably, callus-inducing signals partially restore rhythmicity in *elf3* mutants, revealing feedback from hormonal signaling to the circadian system. These findings identify ELF3 as an integrator of circadian and hormonal inputs that temporally constrains root developmental plasticity.

## Main Text

Lateral root (LR) formation is a major determinant of root system architecture (RSA), influencing plant growth, nutrient uptake, and stress resilience. To sustain RSA, roots generate LRs at regular intervals through a root-specific oscillatory mechanism known as the “root clock” (*1*). This clock controls periodic auxin accumulation in the primary root, producing auxin maxima approximately every six hours at defined “pre-branch sites.” Once these maxima appear in pericycle cell files, auxin-dependent pathways trigger LR initiation, beginning with asymmetric pericycle divisions that establish the lateral root primordium (LRP) (*2*).

Key auxin-responsive transcription factors, including AUXIN/INDOLE-3-ACETIC ACID (AUX/IAA), AUXIN RESPONSE FACTOR (ARF), and LATERAL ORGAN BOUNDARIES DOMAIN (LBD) families, drive these early asymmetric divisions and LRP formation (*3*). Beyond auxin, additional signals such as reactive oxygen species contribute to LR regulation (*4–7*). Later phases of LR development are governed by further molecular complexity. Notably, a recent study revealed that the circadian clock, previously thought to function independently of the root clock, becomes active during late LRP development (*8*). The core circadian clock regulator TIMING OF CAB EXPRESSION 1 (TOC1; also known as PSEUDO-RESPONSE REGULATOR 1 [PRR1]) rephases at this stage and modulates expression of *DIOXYGENASE FOR AUXIN OXIDATION 2* (*DAO2*), an auxin-inactivating enzyme required for proper LR emergence (*8*). These findings indicate that coordinated rhythmic regulation by both the root clock and the circadian clock is essential for successful LR development.

Despite these advances, the mechanisms by which these two timing systems are integrated remain unresolved. In particular, it is unknown at which developmental phase, LR initiation or emergence, the circadian clock interfaces with the root clock to coordinate LR formation. Although circadian rephasing is known to influence later stages of LR development, how circadian rhythms contribute to the initiation of LRP formation remains unclear. A recent study showed that TOC1 regulates *CELL DIVISION CYCLE 6* (*CDC6*), a key driver of the G1–S cell-cycle transition in shoots (*9*), suggesting a direct molecular connection between circadian timing and cell-cycle control. This raises the possibility that circadian regulation may also operate at the onset of LRP initiation.

We tested the hypothesis that the circadian clock may act early in LRP initiation using the callus-inducing medium (CIM) system, a robust platform that reprograms *Arabidopsis* root cells into a pluripotent state and enables controlled induction and analysis of callus formation. CIM, which contains high levels of synthetic auxin (2,4-D) and cytokinin (kinetin), induces callus with strong developmental similarity to early LRPs (*10*). In both petals and leaves, CIM triggers pericycle-like cells to undergo ectopic divisions, accompanied by strong upregulation of core LR regulators (*10*). In roots, CIM causes widespread pericycle activation and vigorous callus formation, with ARF and LBD transcription factors functioning as essential drivers of these early divisions (*10*). Thus, CIM provides a powerful system for dissecting the molecular events underlying LR initiation, allowing amplification and precise analysis of early-stage processes that are otherwise restricted to a few pericycle cells in native LRPs.

### Loss of Circadian Rhythmicity and Emergent TOC1 Waves Couple the Clock to Early LRP initiation

*TOC1* expression is elevated in CIM-induced callus derived from *Arabidopsis* leaves, and *toc1-3* mutants display delayed callus formation (*11*). Other evening-phased components, including *EARLY FLOWERING 3* (*ELF3*) and *ELF4*, are also upregulated, whereas the morning-phased *CIRCADIAN CLOCK ASSOCIATED 1* (*CCA1*) and *LATE ELONGATED HYPOCOTYL* (*LHY*) are repressed, indicating broad circadian gene remodeling during CIM-induced reprogramming (*11*). However, whether CIM alters only the amplitude of clock gene expression or disrupts the period and rhythmicity of the circadian oscillator remained unresolved.

Therefore, we analyzed circadian outputs in CIM-treated roots using *pTOC1::LUC* and *pCCA1::LUC* reporters, widely used to monitor rhythmicity (*12*). Under CIM treatment, *pTOC1::LUC* initially peaked at Zeitgeber Time 16 (ZT16; 16 hrs after the onset of light), similar to controls, but rhythmic oscillations rapidly dampened thereafter, eventually disappearing as expression drifted upward (Fig. 1A). *pCCA1::LUC* showed a similar pattern, with expression suppressed after its first peak followed by a loss of rhythmicity (Fig. 1A). These results show that CIM disrupts the Arabidopsis circadian clock early, eliminating rhythmic expression of both morning (*CCA1*) and evening (*TOC1*) components. Rather than exhibiting a simple phase shift or reduction in amplitude, the clock becomes strongly damped and uncoupled under callus-inducing conditions.

**Figure 1.**
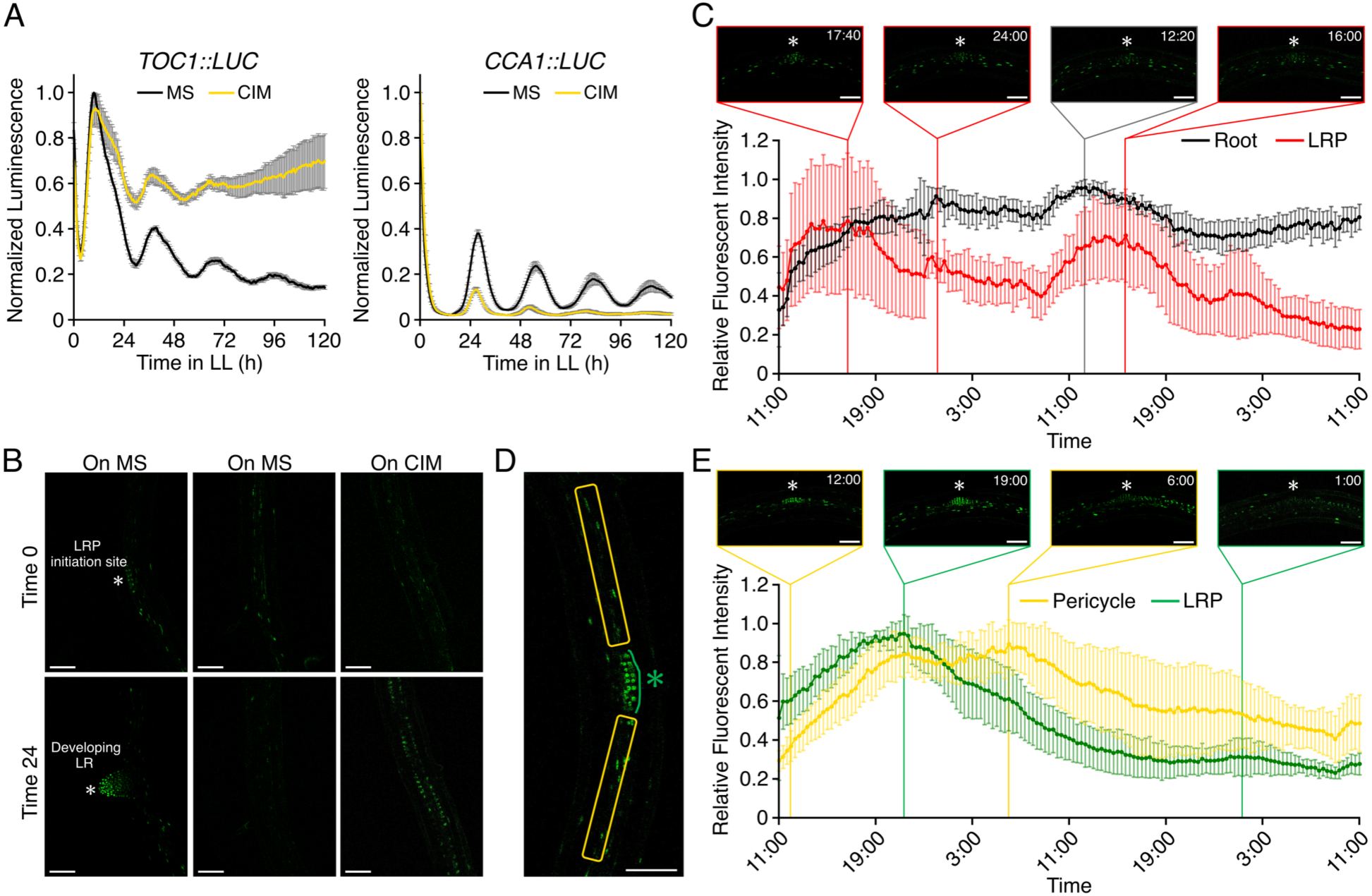
TOC1 showed specific expression patterns in LRP and developing callus induced by CIM. (A) LUC intensity of *pTOC1::LUC* and *pCCA1::LUC* in roots. *pTOC1::LUC* and *pCCA1::LUC* seedlings were grown on MS medium under LD conditions for 6 days, after which roots were excised and transferred to either MS or CIM medium. LUC luminescence intensity was quantified under continuous light condition for 120 hrs. The graph shows the average LUC intensity from 15 samples with SD. Black lines: MS medium, Yellow lines: CIM medium. (B) *pTOC1::TOC1-GFP*/*toc1-2* expression in roots of 7 days old seedlings. Left panels; TOC1-GFP around LRP developing region on MS. Middle panels; TOC1-GFP in the maturation zone on MS, Right panels; TOC1-GFP in the maturation zone on CIM. The images in the upper row represent time 0 after transfer to the respective media, while the images in the lower row represent 24 hrs post-transfer. Scale bars, 75 µm. (C) GFP fluorescent intensities from time-lapse imaging of TOC1-GFP grown on MS. The upper images show root images captured at the indicated time points. White asterisks in the images indicate the position of LRP. The graph in the lower panel shows the relative GFP fluorescence intensity in the entire root of the image (gray line) and the LRP (red line). n = 5, ±SD. (D) The regions quantified for fluorescence intensity in (E). The orange boxes indicate the pericycle cell layers, and the green asterisk indicates the LRP. (E) GFP fluorescence intensity from time-lapse imaging of TOC1-GFP transferred to CIM. The upper images show root images captured at the indicated time points. White asterisks in the images indicate the position of LRP. The graph in the lower panel shows the relative GFP fluorescence intensity in the LRP (green line), pericycle (yellow line), and other cells (gray line) as indicated in (D). n = 5, ±SD. GFP channel images are shown with identical linear display settings (minimum 5,000; maximum 30,000) applied uniformly across all panels for visualization. Quantitative analyses were performed using the unprocessed raw images.

Given the limited cellular resolution luciferase assays, we next examined TOC1 protein dynamics using a *pTOC1::TOC1-GFP* reporter in the *toc1-2/pCCA1::LUC* background. The transgene rescued the *toc1-2* phenotype, confirming its functionality (fig. S1). TOC1-GFP was detected broadly in the root maturation zone and prominently in the meristematic region and early-stage LRPs (Fig. 1B; figs. S2 and S3). After 24 hrs of CIM treatment, TOC1-GFP markedly increased in pericycle cells undergoing division to form callus (Fig. 1B), suggesting that TOC1 participates directly in pericycle reprogramming during LR initiation and callus induction.

Time-lapse imaging further revealed distinct TOC1 dynamics between native LR development and CIM-induced reprogramming. On standard media (Murashige and Skoog; MS), TOC1-GFP oscillated at developing LRP sites (Fig. 1C; fig. S4; Movie S1). In contrast, under CIM treatment, TOC1-GFP peaks showed a temporal offset between LRPs and surrounding pericycle cells (Fig. 1, D and E; fig. S5): fluorescence in LRPs peaked at night and declined, whereas pericycle peaks occurred later and weakened progressively. This wave-like propagation extended both above and below the developing primordium (Movie S5). Strikingly, dividing pericycle cells lacked clear circadian oscillations; instead, TOC1-GFP remained strongly expressed in daughter cells throughout mitosis.

Together, these findings demonstrate that TOC1 expression in pericycle cells does not follow a strict circadian rhythm under CIM but forms spatially propagating expression waves linked to LR initiation. This suggests that TOC1 operates as a dynamic regulator that connects circadian components to pericycle reprogramming during organogenesis. More broadly, the results reveal a mechanistic intersection between the circadian clock and the root clock, indicating that circadian regulators influence the rhythmic gene-expression dynamics underlying both LR initiation and callus formation.

### ELF3 Restrains Root Cell Expansion and Developmental Progression During Callus and LRP Formation

The dynamic changes in TOC1 expression during callus development prompted us to test whether other circadian components contribute to LRP formation. We therefore examined the CIM response of nine core clock mutants (Fig. 2A). Among them, only *elf3-8* produced markedly larger calli than wild-type Col-0 (Fig. 2B). ELF3, a central component of the Evening Complex (with LUX ARRHYTHMO (LUX) and ELF4), restricts downstream gene expression and growth (*13*). Notably, neither *lux* nor *elf4-2* mutants showed increased callus size, and *toc1-2* mutants also resembled the wild type (Fig. 2A). Thus, although TOC1 expression changes accompany pericycle reprogramming, TOC1 itself does not control callus growth. Instead, the data indicate that ELF3 uniquely modulates LR initiation and early development under CIM conditions, prompting us to focus on its function in root reprogramming.

**Figure 2.**
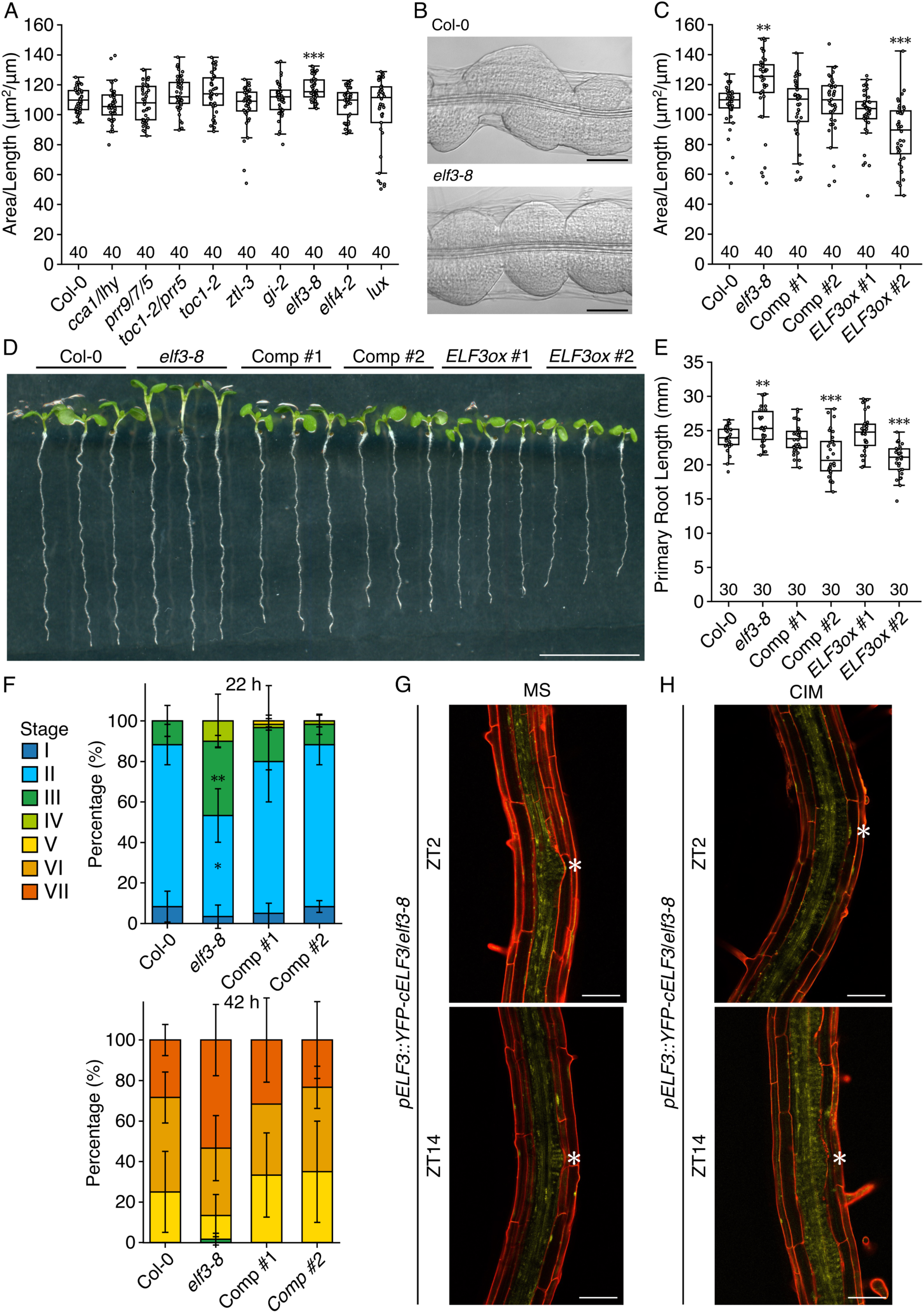
Survey of circadian clock mutants that impact root callus formation. (A) Callus area of plants grown under continuous light for three days after being transferred to CIM following 7 days of growth on MS media under LD conditions (n = 40). \*\*\**p* < 0.001, determined using Student’s *t*-test compared to Col-0 data. (B) Callus images of Col-0 and *elf3-8* grown under the same conditions as (A). Scale bars, 100 µm. (C) Callus area of roots of Col-0, *elf3-8*, two independent *ELF3* complemented lines with *pELF3::YFP-cELF3*, and two independent *ELF3ox* lines treated with CIM for 3 days under continuous light (n = 40). \*\**p* < 0.01, determined using Student’s *t*-test compared to Col-0 data. (D) 7-day-old seedlings of Col-0, *elf3-8*, two *ELF3* complemented lines, and two *ELF3ox* lines. Scale bar, 13 mm. (E) Primely root length of the plants shown in (D) (n = 30). Significant differences from Col-0 were determined using the Student’s *t*-test (\*\*\**p* < 0.001, \*\**p* < 0.01). (F) Percentage of the LRP at different stages of development after 22- and 42-hrs gravistimulation was classified in Col-0, *elf3-8*, and two independent *ELF3* complementation lines. Data are presented as mean ± SE of three biological replicated, with 20 seedlings in each. Significant differences in the number of LRs of each stage from Col-0 were determined using generalized mixed model followed by the Holm’s P-value adjustment in each stage (\*\**p* < 0.01, \**p* < 0.05). Colors on the left denote the different LRP stages. (G) Representative images showing *pELF3::YFP-cELF3* expression in stage III developing LRs of 7-day-old seedlings grown on MS medium, captured at ZT2 and ZT14. Scale bars, 75 µm. (H) Representative images showing *pELF3::YFP-cELF3* expression in stage III developing LRs of 7-day-old seedlings transferred to CIM, imaged at ZT2 and ZT14. Scale bars, 75 µm. Roots were stained with PI solution. Asterisks in the figure indicate LRP position. YFP channel images are displayed using identical linear brightness settings (minimum 2,000; maximum 50,000) applied uniformly across all images to facilitate visualization.

ELF3 is known to regulate circadian timing, flowering, and hypocotyl elongation (*14*). Two *elf3-8* complementation lines and an ELF3 overexpression line rescued the *elf3-8* hypocotyl phenotype (fig. S6) and restored callus size to Col-0 levels (Fig. 2C). We measured cell length in the inner callus layer to assess the developmental basis of the enlarged *elf3-8* calli (fig. S6). *elf3-8* callus cells were significantly longer than those of Col-0, and this defect was rescued in all complementation and overexpression lines (fig. S6). These results demonstrate that enhanced callus development in *elf3-8* arises from increased cell elongation, rather than elevated cell proliferation. Together, these findings identify ELF3 as a key negative regulator of cell expansion during callus formation, providing a mechanistic link between circadian regulation and the cellular growth dynamics underlying pericycle reprogramming and LR initiation.

The enhanced callus phenotype of *elf3-8* suggested that ELF3 broadly regulates root development. We therefore examined primary root growth and LR formation under standard conditions. Seven-day-old *elf3-8* seedlings displayed significantly longer primary roots than Col-0 (Fig. 2D and E). In *elf3-8*, both the meristematic zone length and meristem cell length were increased, whereas the number of meristematic cells remained unchanged (fig. S7). These defects were fully rescued in *ELF3* complementation and overexpression lines, and mature cortical cell length was comparable to Col-0 (fig. S7). Thus, ELF3 restricts primary root growth by limiting cell elongation within the meristem.

This elongation phenotype mirrors the enhanced callus growth observed under CIM treatment. Despite these effects on meristematic growth, *elf3-8* exhibited normal LRP density (fig. S8), indicating that ELF3 does not influence the initiation of LR founder cells. To assess later developmental steps, we monitored LRP progression using gravistimulation. At 22 hrs post gravistimulation (hpg), *elf3-8* LRPs accumulated at stages III–IV, and by 42 hpg, stage VII LRPs were also enriched relative to Col-0. These developmental accelerations were restored in ELF3 complementation lines (Fig. 2F). Therefore, ELF3 does not control LR initiation but acts as a brake on LRP progression, and its loss accelerates advancement through successive developmental stages.

We next analyzed ELF3 expression using *pELF3::YFP-cELF3* in the *elf3-8* background. Although YFP signals were generally weak, ELF3 was broadly detected across the root maturation zone, similar to TOC1-GFP. Within LRPs, YFP-cELF3 accumulated from stage I onward (Fig. 2G; fig. S9). Under CIM treatment, YFP-cELF3 was also strongly expressed in proliferating pericycle cells (Fig. 2H; fig. S10; Movie S3). These results indicate that ELF3 is actively recruited to pericycle cells during both native LR development and CIM-induced proliferation, supporting a model in which loss of circadian periodicity in dividing pericycle cells allows ELF3 to function as a key regulator of LRP developmental pace.

### CIM and ELF3 Rewire Circadian Outputs in Roots

Given that CIM enhances LR formation and that circadian clock mutants can either disrupt or amplify this process, we sought to examine the temporal expression dynamics underlying these responses. Therefore, we generated a high-resolution time course by sampling *Arabidopsis* roots every four hours over a 24-hour period (in duplicate for 12 samples each) following transfer to either CIM or control medium (MS). We focused on the *elf3-8* mutant, a strong loss-of-function allele that exhibits arrhythmic behavior under continuous light and dampened circadian amplitude under light/dark cycles (*15*, *16*), and which displays markedly enhanced callus development under CIM treatment. ELF3 is a central circadian regulator that gates physiological processes to specific times of day, including hypocotyl elongation, and integrates environmental cues such as temperature into the clock (*17*, *18*); loss of ELF3 disrupts these gating functions and results in characteristic developmental phenotypes, including early flowering and constitutive photomorphogenesis (*19*). To dissect how hormonal signals and circadian regulation interact to shape gene expression dynamics during root developmental reprogramming we analyzed roots from both Col-0 and *elf3-8* grown under MS or CIM conditions, yielding four complete time courses: Col-0 on MS, Col-0 on CIM, *elf3-8* on MS, and *elf3-8* on CIM (Fig. 3A).

**Figure 3.**
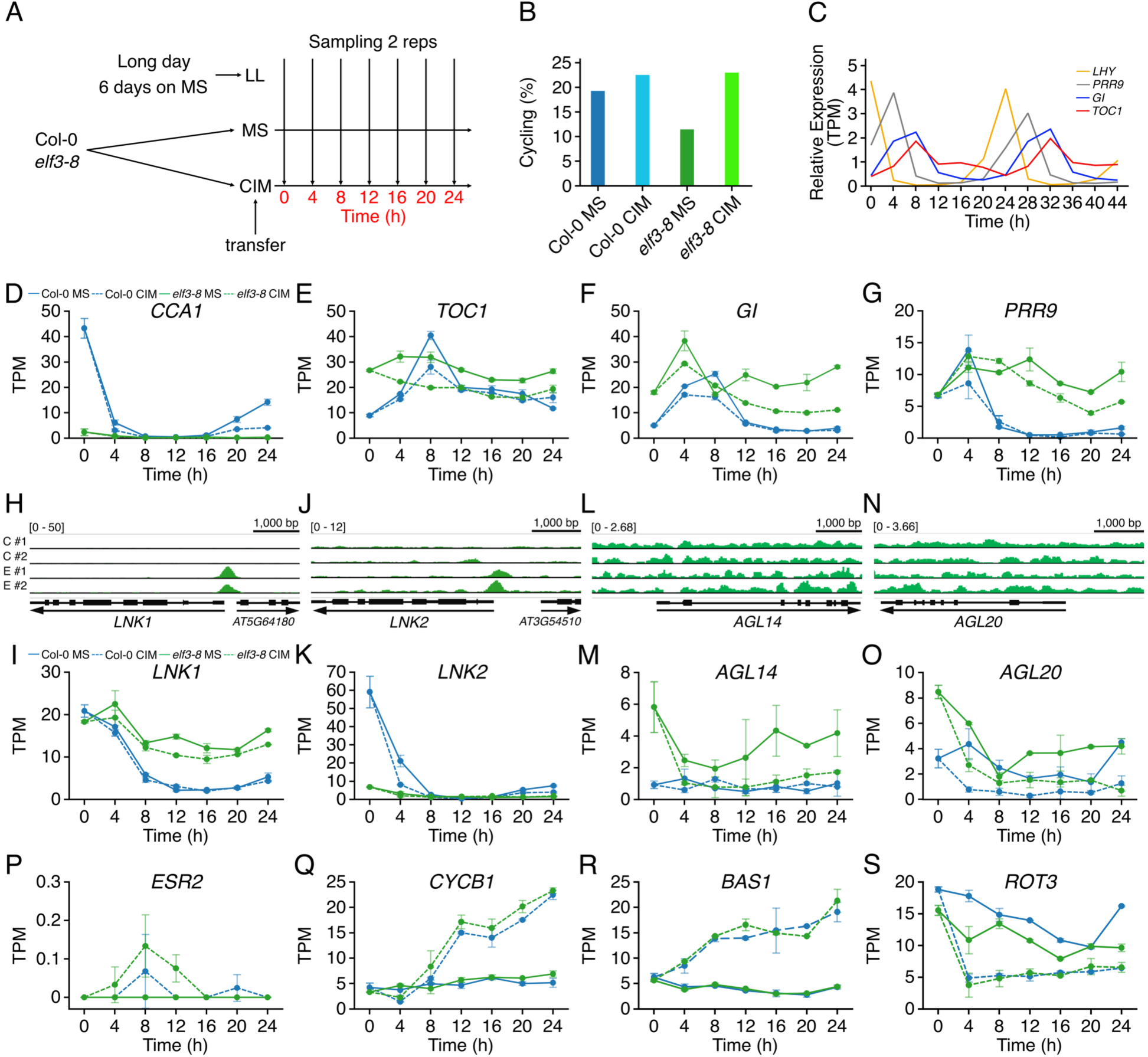
*ELF3* gates CIM responsiveness. (A) *Arabidopsis* root time course experimental design. Wild type Col-0 and *elf3-8* mutant plants were grown under long days for six days on MS media and then transferred to continuous light (LL) and CIM treatment. Samples were taken in duplicate every four hours for Col-0 CIM, Col-0 MS, *elf3-8* CIM and *elf3-8* MS. (B) Percent of cycling genes in *Arabidopsis* roots. (C) Core circadian clock genes cycle with the expected relative phase; *LHY* (yellow) peaks at dawn (Circadian Time 0; CT0); *PRR9* (grey) peaks at CT4; *TOC1* (orange) peaks at CT8; and *GI* (blue) peaks at CT8. (D-G) The *elf3-8* mutant with and without CIM result in expression damping in CCA1 (D), a phase advance in TOC1 (E) and *GI* (F), and a phase delay in *PRR9* (G). (H-O) ELF3 binds to the promoter regions of *LNK1* and *LNK2* and negatively regulates *LNK1* expression. Although ELF3 also binds to the *LNK2* promoter, it does not appear to be involved in regulating *LNK2* expression. In addition, ELF3 does not bind to the *AGL14* and *AGL20* promoter, whose expression is derepressed in the *elf3-8*. (H, J, L, and N) Visualization of ELF3-FLAG ChIP-seq data using GVI. “Black boxes” indicate ORFs (thick boxes: exon; lines: intron) and black arrows beneath each ORF represents the direction of transcription. The scale for peak detection on the Y-axis is shown to the left of each graph. The scale bars above graphs indicate 1,000 bp. C#1; Col-0 biorep 1, C#2; Col-0 biorep 2, E#1; ELF3-FLAG biorep 1, E#2; ELF3-FLAG biorep 2(I, K, M, and O) Gene expression patterns of *LNK1*, *LNK2*, *AGL14*, and *AGL20* extracted from RNA-seq data. (P-S) The expression patterns of representative genes exhibiting rhythmic expression in CIM. Data were extracted from the time course RNA-seq data set. All genes expression data are shown with blue representing Col-0 and green representing *elf3-8*; solid lines indicated MS medium, and dashed lines indicate CIM treatment. The X-axis indicates treatment time (hrs), and the y-axis represents expression values with transcripts per million (TPM).

These datasets were analyzed for rhythmicity, phase and period (*20*) (Data S1). Across the time courses, approximately 70% of expressed genes were detected in each condition and Col-0 plants on MS had 20% genes cycling consistent with similar above ground experiments (*21*) (Fig. 3B). In contrast, only 11% of genes cycled on MS in the *elf3-8* mutant background, which suggests *elf3-8* is not arrhythmic in roots or previous studies in shoots missed cycling genes since they only looked at marker genes (*15*). Interestingly, we observed a modest increase (∼3%) in the number of cycling genes under CIM compared to MS in the Col-0 as well as cycling being restored in *elf3-8* under CIM (∼20% cycling vs. 11%) (Fig. 3B). This observation implies that the hormonal cues induced by CIM, specifically the synthetic auxin (2,4-D) and cytokinin, may interact with the circadian system to modulate rhythmic gene expression in a way that primes the root for developmental reprogramming, such as LR initiation and callus development.

The ability of CIM to restore rhythmicity in *elf3-8* further supports the hypothesis that *ELF3* modulates hormone-responsive gene networks through circadian gating, similar to its established role in the shoot. Gene Ontology (GO) enrichment analysis of the genes whose rhythmic expression was restored in *elf3-8* under CIM revealed overrepresentation of responses to karrikin (GO:0080167), UV-B light (GO:0010224), and red/far-red light (GO:0009639) (Data S2). Notably, this gene set includes *HY5* and *HYH*, transcription factors previously implicated in auxin-mediated root growth (*22*), suggesting that *ELF3* may indirectly regulate these hormone-and light-responsive genes via circadian mechanisms. In aerial tissues, *ELF3* is known to restrict the activity of transcription factors such as *PIF4* to specific times of day, thereby coordinating light, hormone, and circadian signals to regulate growth (*17*). By analogy, we propose that *ELF3* similarly modulates root growth by acting through *HY5* and *HYH*, extending a conserved temporal gating mechanism from shoot to root development.

Consistent with this hypothesis *hy5* and the *hy5/hyh* double mutant, but not the *hyh* single mutant result in reduced callus area (fig. S11). We further analyzed LR development in response to gravistimulation in *hy5*, *hyh*, and *hy5*/*hyh* mutants (fig. S11). At 22 hpg, the proportion of stage I LRP was higher in *hy5* and *hy5*/*hyh* than in Col-0. At 42 hpg, stage II to IV LRP were more abundant in *hy5* and *hy5*/*hyh* compared to Col-0. Consistent with the reduced callus area upon CIM treatment observed in *hy5* and *hy5*/*hyh*, LR development was delayed in these mutants. Our findings support a parallel function for *ELF3* in roots, where it may regulate temporal aspects of hormone signaling and developmental transitions, potentially through transcriptional regulators like *HY5*, positioning ELF3 as a central temporal gatekeeper in both shoot and root development.

Next, we validated our time-course data and assessed whether CIM influences circadian clock function by examining the expression patterns of 91 well-characterized clock, flowering and light signaling genes (*20*, *23*) (Fig. 3C). In Col-0 roots on MS, 36% of the clock, flowering and light signaling genes exhibited rhythmic expression, far less than previous findings in whole seedlings, where ∼60% of clock genes were reported to cycle. In contrast, the *elf3-8* mutant showed a substantial reduction in the number of cycling clock genes, with only 12% maintaining rhythmic expression (Data S3). This mirrors the global dampening of rhythmicity observed in *elf3-8* and supports the known role of *ELF3* in sustaining circadian rhythms (Fig. 3B). However, the persistence of some rhythmic genes indicates that *elf3-8* is not entirely arrhythmic in root tissues (Data S3). In Col-0, CIM reduced the proportion of cycling clock genes from 36% to 25%, suggesting that CIM can dampen circadian rhythmicity in roots. Conversely, in *elf3-8*, CIM increased the number of cycling clock genes from 12% to 32%, indicating a partial restoration of rhythmicity in the absence of functional *ELF3* (Fig. 3B). These results suggest that *ELF3* acts as a negative regulator of CIM-induced LR development, potentially by suppressing TOD-dependent transcriptional programs. The ability of CIM to partially rescue rhythmic expression in *elf3-8* implies crosstalk between hormone signaling and circadian networks, where developmental cues may compensate for defects in core clock components.

Since CIM treatment led to a reduction in the number of cycling core clock genes, we next asked whether CIM directly disrupts the circadian clock or acts downstream of it. To test this, we examined the rhythmic expression of key circadian clock components (*CCA1, LHY, TOC1/PRR1, PRR3, PRR5, PRR7, PRR9, GIGANTEA*[*GI*]*, LUX, and ELF4*) in Col-0 roots

under both MS and CIM conditions. In Col-0 roots, CIM did not significantly disrupt the rhythmicity or phase of core clock gene expression (Fig. 3D-G), suggesting that CIM acts downstream of the circadian oscillator rather than perturbing its core function. In contrast, *elf3-8* roots displayed specific and notable disruptions in core clock gene expression. *CCA1* and *LHY* expression levels were markedly dampened, while *PRR1*, *GI*, *PRR3*, and *PRR9* exhibited mildly elevated expression compared to Col-0 (Fig. 3D-G; fig. S12). These changes are consistent with the known role of *ELF3* in modulating amplitude and phase within the circadian system, particularly in gating inputs and outputs of the clock. Moreover, CIM further altered the expression patterns of core clock genes in the *elf3-8* background, indicating that *ELF3* is necessary for buffering or gating hormone-induced transcriptional changes. Since these genes remain largely unaffected by CIM in Col-0, the enhanced sensitivity in *elf3-8* suggests that *ELF3* plays a key role in mediating cross-talk between the circadian system and hormone signaling pathways. While *ELF3* itself cycles with the correct phase in Col-0, its expression is modulated by CIM treatment. This responsiveness further supports the idea that *ELF3* functions as a gating node, regulating the temporal integration of circadian and hormonal signals. These results indicate that the circadian clock operates upstream of hormone-induced developmental responses, and that *ELF3* acts as a critical mediator at this interface, consistent with its role as a central hub in the EC.

Although *ELF3* lacks intrinsic DNA-binding activity, it functions as a transcriptional repressor through its role in the EC, where it forms a regulatory module with *ELF4* and *LUX*. Previous ChIP-seq studies in *Arabidopsis* have identified ELF3-associated genomic regions, supporting its role in indirectly targeting DNA via complex formation (*25*). We performed ChIP-seq using roots of *35S::ELF3-FLAG*/*elf3-8* plants treated with CIM for 24 hrs to investigate ELF3 binding in the root under callus-inducing conditions. We identified 44 significant *ELF3*-associated peaks located within 3,000 bp upstream of gene coding regions (Fig. 3H-O; Data S4). These included multiple core circadian clock genes such as *CCA1*, *PRR9*, *PRR7*, *LUX*, *GI*, *LNK1*, *LNK2*, *LNK3*, and *LNK4* (fig. S13). Consistent with these binding events, expression patterns of several clock genes were altered in *elf3-8* compared to Col-0 (fig. S12). Specifically, *CCA1*, *PRR9*, and *PRR7* exhibited a 2-4 hr phase delay, while *GI* and the *LNK* family genes showed elevated expression levels during the night, when these transcripts are normally suppressed in Col-0 plants (Fig. 3F, I, and K). These findings support the idea that *ELF3* represses key transcription factors in the circadian clock, helping to establish proper timing and amplitude of clock gene expression. The loss of *ELF3* disrupts this tightly coordinated regulatory network, leading to widespread misregulation of rhythmic gene expression. The exaggerated callus phenotype observed in *elf3-8* under CIM is likely one such output of this dysregulated circadian system, reflecting the broader impact of clock dysfunction on developmental reprogramming.

We identified a second class of genes that were responsive to CIM treatment independently of the ELF3 genetic background (Fig. 3P-S). These genes do not exhibit rhythmic expression under MS conditions in either Col-0 or *elf3-8*, yet they show a coordinated and robust transcriptional response following CIM exposure regardless of genotype. Despite their lack of cycling under control conditions, these genes appear to be highly regulated in response to developmental reprogramming, likely through hormonal cues activated by CIM. Functionally, this gene set shares enrichment for pathways involved in hormone responsiveness (auxin-centric), developmental timing and morphogenesis, metabolic and energetic coordination, and transport and cell wall execution, with six overrepresented terms associated with root development (Data S5). The consistency of their expression responses across genotypes suggests these genes operate downstream of ELF3 and, more broadly, the core circadian clock, given that they are not regulated under MS and do not cycle in the absence of CIM. Notably, their temporal expression patterns under CIM suggest that their transcription is still gated by the circadian system. For example, even though they do not cycle under MS, the timing of their induction or repression under CIM follows predictable circadian phases consistent with gating. These genes can be broadly categorized into at least four temporal expression classes: genes that decrease at dawn; genes that increase in the early morning and decline; genes that peak around midday and taper off; genes that show a continuous increase across the day; and genes that decrease rapidly (Fig. 3P-S). These distinct expression profiles suggest that this class of genes represents hormone-responsive targets whose activation is modulated by circadian phase. Moreover, their responsiveness in both Col-0 and *elf3-8* backgrounds indicates that, while *ELF3* influences many clock-regulated processes, CIM-induced hormonal pathways can partially bypass or override *ELF3* function, possibly through circadian gating mechanisms that are *ELF3*-independent.

Finally, we identified a distinct class of genes that respond to both CIM treatment and ELF3 activity, which may represent ELF3-regulated effectors involved in LR development. These genes are repressed in the *elf3-8* mutant but are induced or modulated by CIM, suggesting they function downstream of ELF3 while remaining responsive to hormone-induced developmental reprogramming (Fig. 3M and O). Notably, this group includes several MADS-box transcription factors from the AGAMOUS-like (AGL) family, which are known to regulate diverse developmental processes. Within this group, *AGL20*/*SOC1*, *AGL14*/*XAL2*, and *AGL19* show this dual regulation pattern, while other family members like *AGL12*/*XAL1* exhibit expression more typical of CIM-responsive but ELF3-independent genes. *AGL20*/*SOC1* is classically known as a key flowering-time regulator, but recent work has also shown that it modulates root meristem activity, including cell division and differentiation, thus influencing root architecture (*26*). The dual responsiveness of *AGL20*/*SOC1*to ELF3 and CIM suggests it may help coordinate LR initiation by integrating circadian and hormonal signals. AGL14/XAL2 is essential for auxin transport and distribution in the root and plays a central role in root formation and maintenance. It regulates PIN gene expression, controlling auxin efflux and directional transport, critical steps for LR patterning (*27*). Supporting this, RNA-seq data from 7-day-old *xal2-2* roots revealed misregulation of genes associated with cell wall remodeling and LR initiation, many of which also respond to CIM, suggesting that XAL2 operates downstream of ELF3 and hormone pathways (*26*). AGL12/XAL1 is implicated in cell cycle regulation, another key step in LR primordia formation. Its expression profile under ELF3 and CIM supports a role in mediating hormone-circadian coordination of cell cycle reactivation during reprogramming. Collectively, these findings suggest that ELF3 promotes LR initiation not only through circadian control and hormonal gating but also by regulating a transcriptional network of MADS-box genes that act as developmental integrators, processing temporal, hormonal, and spatial signals to fine-tune root system architecture. However, our ChIP-seq analysis revealed that these AGLs were not direct targets of ELF3 (Fig. 3L and N).

### The ELF3-LNK-AGL Module Control Callus Growth and LRP Tempo

We next examined direct ELF3 targets to identify downstream effectors through which ELF3 regulates LR reprogramming. All four *LNK* genes, known EC targets, were misregulated in *elf3-8* regardless of CIM treatment (Fig. 3I and K; Data S1), indicating that ELF3 normally constrains their expression. Consistent with this, the *lnk1/lnk2* double mutant generated significantly larger calli under CIM, and this phenotype was rescued by *pLNK1::LNK1-FLAG* (Fig. 4A and B).

**Figure 4.**
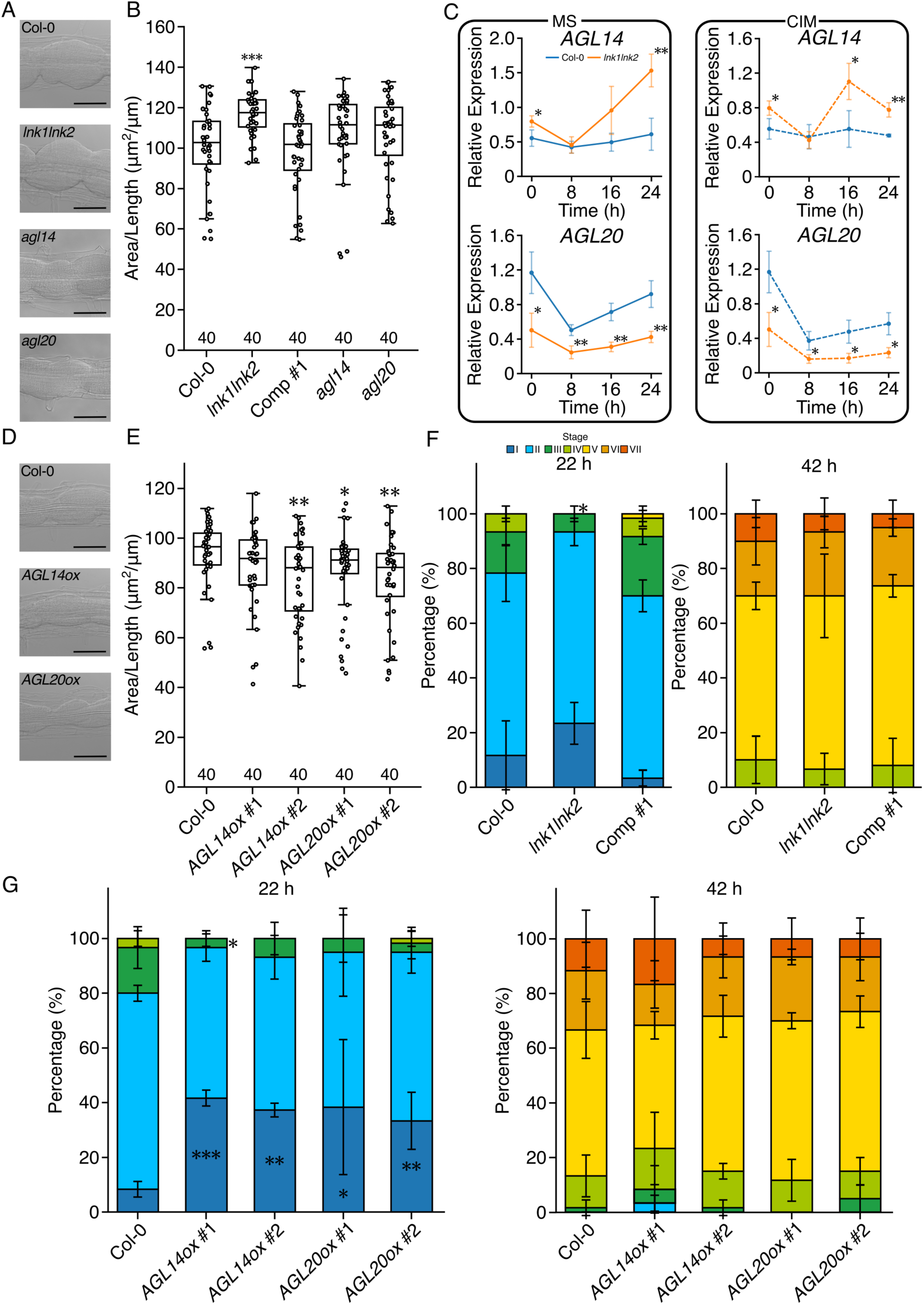
ELF3 downstream genes involved in LR development. (A) Callus images of Col-0, *lnk1*/*lnk2*, *agl14*, and *agl20* grown under continuous light for three days after being transferred to CIM following 7 days of growth on MS media under LD conditions. Scale bars, 100 µm. (B) Callus area of the roots shown in (A) (n = 40). \*\*\**p* < 0.001 determined using Student’s *t*-test compared to Col-0 data. (C) qPCR analysis of ELF3 downstream genes in roots of Col-0 and *lnk1*/*lnk2* treated with CIM for 0, 8, 16 and 24 hrs (n = 3, ±SD). Significant differences from Col-0 were determined using the Student’s *t*-test at each time point (\*\**p* < 0.01, \**p* < 0.05). Genes expression data are shown with blue representing Col-0 and orange representing *lnk1*/*lnk2*; solid lines indicated MS medium, and dashed lines indicate CIM treatment. (D) Callus images of Col-0, *AGL14ox*, and *AGL20ox* grown under continuous light for three days after being transferred to CIM following 7 days of growth on MS media under LD conditions. Scale bars, 100 µm. (E) Callus area of the roots shown in (D) (n = 40). \**p* < 0.05 and \*\**p* < 0.01 determined using Student’s *t*-test compared to Col-0 data. (F and G) Percentage of the LRP at different developmental stages were counted after 22- and 42-hrs of gravistimulation in Col-0, *lnk1*/*lnk2*, and *LNK1* complemented line (F); and Col-0 and *AGLox* lines (G). The roots in (G) were treated with 5 µM estradiol for inducing *AGL* genes. Data represent mean ± SE from three independent biological replicates, each consisting of 20 seedlings. Statistical significance relative to Col-0 at each stage was determined using a generalized mixed model followed by Holm’s P-value adjustment in each stage (\*\**p* < 0.01, \**p* < 0.05). Colors on the top denote the different LRP stages.

LNK1/LNK2 act as transcriptional coactivators that activate clock genes such as *PRR5* and *TOC1* via RECEILLE (RVE) factors and are essential for proper circadian period and light signaling (*28*). Together, these findings indicate that LNK1/LNK2 restrain callus proliferation, and that their repression by ELF3 contributes to the enhanced reprogramming observed in *elf3-8*. Misregulation of LNK proteins may thus disturb circadian gating, linking ELF3 function to both developmental timing and cellular plasticity.

Previous transcriptomic studies reported reduced *AGL20* expression in *lnk1*/*lnk2* (*28*). Similarly, *AGL20* expression was lower in the *lnk1*/*lnk2* than in the wild type under both MS and CIM treatment (Fig. 4C). In contrast, *AGL14* expression was significantly elevated in *lnk1/lnk2* at multiple time points under CIM and even on MS medium (Fig. 4C). Given the documented reciprocal regulation between *AGL14* and *AGL20* (*26*, *29*), the concomitant upregulation of *AGL14* and reduction of *AGL20* supports the existence of a feedback mechanism governing MADS-box gene dynamics. Functionally, *AGL14ox* and *AGL20ox* lines exhibited strongly reduced callus formation (Fig. 4D and E), whereas *agl14* and *agl20* single mutants were indistinguishable from Col-0 (Fig. 4A and B), indicating functional redundancy in restricting proliferation. Together, these findings demonstrate that overexpression of specific MADS-box genes is sufficient to suppress callus proliferation and that LNK1 is required to maintain appropriate *AGL14* and *AGL20* expression patterns during developmental reprogramming, positioning the ELF3–LNK module as a key regulator linking circadian signaling to pericycle plasticity.

We performed a gravistimulation assay to test whether the ELF3–LNK1–AGL axis also influences LRP development. Although *lnk1/lnk2* mutants produced enhanced calli, their LRP pattern differed from *elf3-8*. At 22 hpg, *lnk1/lnk2* roots displayed a reduced proportion of stage III LRPs compared with Col-0, but by 42 hpg their stage distribution had largely normalized (Fig. 4F). In contrast, *AGL14ox* and *AGL20ox* lines, both impaired in callus formation, accumulated stage I LRPs at 22 hpg and recovered by 42 hpg, indicating a delay in early LRP progression (Fig. 4G). Consistent with their weak callus phenotypes, *agl14* and *agl20* mutants resembled Col-0 in LRP stage distribution (fig. S14).

Closer comparison revealed that LRP progression is accelerated in *elf3-8* but transiently delayed in *lnk1*/*lnk2*, despite both showing enhanced callus proliferation. If callus and LR development were mechanistically coupled, these phenotypes would appear contradictory. Instead, our data suggest that callus enhancement reflects a common consequence of disrupted circadian gating, whereas LRP tempo depends on the interaction with parallel developmental and hormonal pathways. In *lnk1/lnk2*, loss of rhythmic gating may shift the temporal “window” for LR initiation, causing a brief delay. In *elf3-8*, although LNK rhythmicity is absent, overactivation of compensatory pathways, likely auxin- or hormone-responsive, accelerates LRP progression.

Together, these findings show that the LNK1–AGL module controls both callus development and the developmental tempo of LRPs. Loss of *LNK1*/*LNK2* causes a mild, recoverable delay, whereas sustained misexpression of *AGL14*/*AGL20* imposes a stronger bottleneck on primordia advancement. These results support a model in which ELF3, through repression of *LNK1* and downstream *AGL*s, integrates circadian and hormonal cues to coordinate callus and LRP development.

Taken together with the accelerated LRP development observed in *elf3-8* mutants, our findings support a model in which ELF3 functions as a central temporal integrator, coordinating callus formation and LR development through transcriptional regulators such as HY5, LNK1, and MADS-box proteins including AGL14 and AGL20. These downstream factors integrate circadian timing cues with hormonal signals to fine-tune root developmental transitions.

Importantly, our time-lapse imaging of TOC1-GFP revealed that circadian oscillations are dampened in dividing pericycle cells and replaced by spatial “waves” of expression, suggesting that the circadian clock and the root clock are overlapping rather than independent oscillatory systems. We propose that ELF3 maintains this integration by gating hormone-responsive transcriptional modules and stabilizing rhythmic outputs, thereby coupling circadian periodicity to the spatial patterning activities of the root clock. This model advances a broader hypothesis: that the circadian system allocates energy and assimilates aboveground metabolic status, which is then transmitted via ELF3 to root tissues to ensure that growth and regenerative reprogramming occur only when sufficient or specific resources are available. Given prior evidence that ELF3 can mediate shoot-to-root communication, our results suggest that ELF3 links circadian control of energy distribution with local developmental decisions in the root. By positioning ELF3 at the nexus of circadian timing, metabolic status, and hormone signaling, we uncovered a previously uncharacterized regulatory framework that unites the circadian and root clocks and propose that this integration represents a fundamental mechanism by which plants coordinate whole-organism energy balance with the timing and spatial dynamics of organogenesis.

## Supporting information

Supplemental informations

## Funding

This work was supported by JSPS KAKENHI Grant number 24K02049 (to H.T.).

## Author contributions

Conceptualization: T.M., H.T.; Funding acquisition: H.T.; Investigation: S.N., S.U., A.M., S.K., K.M., K.A., T.S., S.I., S.S., N.M., T.M., H.T.,; Data analysis: A.M., K.M., T.S., S.I., N.N., T.M., H.T.; Methodology: T.M., S.I., N.N., T.M., H.T.; Visualization: N.S., S.U., S.K., A.K., H.T.; Writing - original draft: S.N., N.N., T.M., H.T.

## Competing interests

Authors declare that they have no competing interests.

## Data and materials availability

All data are available in the main text or the supplementary materials. RNA-sequencing raw sequence reads have been deposited in DDBJ DRA (https://www.ddbj.nig.ac.jp/index-e.html) with the accession number PRJDB40024. ChIP-sequencing raw sequence reads have been deposited in GEO with the accession number GSE316052.

## Supplementary Materials

Materials and Methods

Figs. S1 to S14

Table S1

References (30–54)

Movies S1 to S3

Data S1 to S5

## Notes

### Competing Interest Statement

The authors have declared no competing interest.

