## Supplemental informations for "The circadian clock gates lateral root development"

**The PDF file includes:**

Materials and Methods
Figs. S1 to S14
Table S1
References (30-51)

**Other Supplementary Materials for this manuscript include the following:**

Movies S1 to S3 (.mp4)
Data S1 to S5 (.xlsx)

### Materials and Methods

#### Plant materials and growth conditions

*Arabidopsis* Col-0 was used as the wild type. Seeds of *CCA1::LUC*/Col-0 (*30*), *TOC1::LUC*/Col-0 (*12*), *ztl-3* (*12*), *elf3-8* (*16*), *cca1*/*lhy* (*31*), *prr9/7/5* (*30*), *toc1*/*prr5* (*32*), *toc1-2* (*32*), *gi-2*, *elf4-2* (*33*), *lux-cr* and *lnk1*/*lnk2* (*31*) were used in this study. *hy5*, *hyh*, and *hy5*/*hyh* mutant seeds were kindly provided by Tomonao Matsushita (Kyoto university, Japan). *agl14* (CS869195) and *agl20* (SALK_138141C) were obtained from the SALK collection and the seed stock center of the Arabidopsis Biological Resource Center. These mutants were genotyped using left-border primers on the T-DNA, right-side primers on the genome, and left-side primers on the genome (Table S1).

All seeds were sterilized with 1% bleach and 0.05% Triton X-100 for 5 min and washed three times with sterilized water. Seeds germinated on Murashige and Skoog (FUJIFILM Wako pure chemical) medium supplemented with 1% sucrose and 1% agarose after 2 days at 4˚C. Plants were grown vertically in a chamber (Panasonic) at 22˚C with 16 hrs light and 8 hrs dark conditions. For callus induction, 7-day-old seedlings were transferred onto callus-inducing medium (CIM; Murashige and Skoog agarose plates containing 0.5 mg/ml 2,4-D (Sigma aldrich) and 0.05 mg/ml kinetin (Sigma aldrich)) at Zeitgeber Time (ZT) 2, then plates were placed in continuous light condition.

#### Plasmid construction and plant transformation

Genomic DNA from Col-0 was used as a template for the amplification of 2,000 bp upstream region of *pELF3* for promoter cloning and from 1,882 bp upstream of translation start site to 2,634 bp downstream region of *pTOC1::TOC1*. One base of 5' dA overhang was added to the PCR amplicon of *pELF3* using Taq polymerase (Takara Bio Inc., Shiga, Japan), which was then cloned into pENTR5'-TOPO (Thermo Fischer Scientific, Waltham, MA, USA) and named pENTR5'-*pELF3*. For *pTOC1::TOC1* cloning, *pTOC1::TOC1* PCR amplicon was cloned into the pENTR/D-TOPO by In-Fusion reaction (Takara Bio inc.) and named pENTR-*pTOC1::TOC1*.

For *pLNK1-LNK1-FLAG* cloning, the promoter and coding region of *LNK1* was amplified from Col-0 genome and cloned into the pBA002a-PF5-FLAG binary bector, which is a derivative of pBA0002a (*35*) binary vector (*31*). For *35S::ELF3-FLAG* cloning, *the ELF3* coding region was amplified from Col-0 cDNA pool, and sub-cloned into pBS-FLAG (*35*). The resulting *ELF3* and 3 x FLAG region were amplified by PCR assembled and cloned into binary vector pSK1. For YFP-*ELF3*, *AGL14-YFP* and *AGL20-YFP* cloning, cDNA regions were amplified via PCR using the Col-0 root cDNA library as a template, and the amplicons were inserted into 3' end (for *YFP-ELF3*) and 5' end (for *AGL14-YFP* and *AGL20-YFP*) of the pDONR201-*YFP* plasmid (*36*) using the NEBuilder HiFi DNA Assembly Cloning kit (New England Biolabs).

For the *pELF3::YFP-cELF3* construct, pENTR5'-*pELF3* and *YFP-cELF3* containing pDONR201 were cloned into R4pGWB550 (*37*) using LR Clonase II (Thermo Fischer Scientific). For the *pTOC1::TOC1-GFP*, pENTR-*pTOC1::TOC1* was recombined into the pBA-PF5-GW-GFP destination vector (*38*) using LR clonase II (Thermo Fischer Scientific). For the *pXVE::AGL14-YFP* and *pXVE::AGL20-YFP*, *AGL14-YFP* or *AGL20-YFP* containing pDONR201s were cloned into pMDC7 (*39*) using LR Clonase II. The resulting plasmids (*pELF3::YFP-ELF3*, *pTOC1::TOC1-GFP*, *pBA-pLNK1-LNK1-FLAG*, *pSK1-ELF3-FLAG*, *pXVE::AGL14-YFP*, and *pXVE::AGL20-YFP*) were transferred into *Agrobacterium tumefaciens*

(C58C1 pMP90) cells and transformed into *toc1-2/CCA1::LUC*, *elf3-8*, *lnk1/lnk2* (*31*), and Col-0. *pLNK1-LNK1-FLAG* construct compliments long hypocotyl phenotype of *lnk1/lnk2*.
To make *lux-cr*, DNAs corresponding to two guide RNAs for CRISPR-Cas9-mediated genome editing of LUX were cloned into p*KIR1.1* (*40*). The resulting *pKIR1.1-LUX* was transformed into *CCA1:LUC*. The selected *lux-cr* mutant has 20 bp deletion (352 to 371 nt 3' side of translation start site), producing the premature stop codon. The *lux-cr* mutants show long hypocotyl, similar to other *lux* mutants (*41*). All primers used for cloning were listed in Table S1.

**Luminescence-based circadian rhythm assays**
A luminescence-based circadian rhythm assay was performed as previously described (*32*). Assays were performed in the growth chamber under constant white light conditions and at 22°C. Period lengths were determined as previously demonstrated (*32*).

**Phenotypic and microscopic analysis**
To measure hypocotyls and whole root length, the seedlings were scanned using a flatbed scanner GT-7400U (Epson) while growing on plates. Hypocotyl and root length were measured by importing the scanner images into ImageJ image-processing software (imagej.net/software/fiji/).
The newly formed callus area derived from pericycle of the root maturation zone was measured with cleared roots after 3 d of incubation on CIM. For root clearing, roots were directly mounted on the slide glasses with clearing solution (a mixture of 20 g of chloral hydrate in 1 ml glycerol and 6 ml water). Then the callus images were obtained using a Leica DMI 6000B-AFC (Leica Camera) and measured callus area by imageJ software.
For callus cell length measurement, plant roots incubated on CIM were fixed and cleared with ClearSee$^{TM}$ solution (FUJIFILM Wako pure chemicals) as described previously (*42*). Briefly, plant roots fixed with 4% (w/v) paraformaldehyde (PFA) and 0.01% Triton in PBS under vacuum (0.04 MPa) for 1 hr at room temperature. Fixed samples were washed twice in PBS and cleared with ClearSee$^{TM}$ at room temperature. For complete clearing, the solution was replaced flesh ClearSee$^{TM}$ solution every three days for 9 days at room temperature. For cell wall staining, cleared roots were immersed in 0.1% (w/v) Fluorescent Brightener 28 (FUJIFILM Wako pure chemicals) in ClearSee$^{TM}$ solution for 24 hrs at dark. The samples were washed twice for 30 min each in ClearSee$^{TM}$ solution at room temperature in the dark. Cell wall stained samples were imaged using a Leica SP8 system (Leica) with 405 nm excitation and 425-475 nm emission. Obtained images were used for measurement of callus cell length by imageJ software.
For time-lapse imaging, a Lab-Tek Chambered Coverglass w/cvr (Thermo Fisher Scientific) was used as described previously (*43*). The chamber containing the plants were imaged with a confocal microscopy Leica SP8 system with 488 nm excitation and 500-550 nm emission for GFP (for *pTOC1::TOC1-GFP/toc1-2* and *pELF3::YFP-cELF3/elf3-8*). Time-lapse images were taken by the LAS X every 20 min for 48 hrs under continuous light condition. Assembling of images were also done with LAS X.
Quantifying the GFP intensity of confocal images was based on previously described methods (*43*). The confocal images were analyzed with imageJ by tracing the outlines of each cell files or tissues (lateral root primordium, pericycles, and entire roots integrated cells of endodermis, cortex and epidermis). The means of the signal were quantified with the "Plot Profile" option.
LR numbers (from stage I to VII [emerged LRP]) in the whole roots of 10-day-old seedlings (7 days under 16 hrs light and 8 hrs dark condition then transfer continuous light condition for 3

days) were counted using a Leica DMI 6000B-AFC (Leica). Stage classification was performed as previously described by Malamy and Benfey (*44*). Roots were cleared using a clearing solution.
For LR induction by gravistimulation, vertically grown 4-day-old seedlings under 16 hrs light and 8 hrs dark condition then transfer continuous light condition for 1 days were rotated 90˚. LRP stages were analyzed after 22- and 42-hrs gravistimulation using a Leica DMI 6000B-AFC (Leica). Stage classification was performed as previously described by Malamy and Benfey (*44*). Roots were cleared using a clearing solution. Experiments were repeated independently three times (n = 20 in each experiment).

**Image processing**
Raw image data were acquired using a Leica LAS X system and recorded as 16-bit images. All images were imported into ImageJ for processing. For visualization purpose, the display range of each channel was adjusted using *Image* > *Adjust* > *Brightness*/*Contrast* by setting the minimum and maximum displayed values as specified below. These adjustments affected only image display and did not modify the underlying pixel intensity values. All quantitative analysis were performed using the unprocessed raw images.
Lookup table (LUTs) were applied as follows: propidium iodide (PI) staining was displayed in red, SCRI staining in grayscale, GFP in green, and YFP in yellow. Individual channels were then merged to generate composite images.
To improve visualization of TOC1-GFP and YFP-ELF3 expression patterns, brightness adjustments were applied uniformly and linearly to the confocal GFP and YFP channel images using ImageJ. For TOC1-GFP, identical brightness settings were applied to all images, and no nonlinear, selective, or local modifications were performed. The same uniform linear adjustment was applied to all YFP-ELF3 images. These adjustments were made solely to enhance visualization and did not alter the relative intensity relationships or affect any quantitative interpretation of the data.
For TOC1-GFP signals, the minimum and maximum displayed values were set to 5,000 and 30,000, respectively, for Fig. 1B-E, fig. S4, and fig. S5; to 0 and 35,000 for fig. S2; and to 0 and 30,000 for fig. S3
For YFP-ELF3 signals, the minimum and maximum displayed values were set to 2,000 and 50,000 for Fig. 2G and H, and to 0 and 35,000 for fig. S9 and fig. S10.
For PI staining in fig. S10, the minimum and maximum displayed values were set to 0 and 10,000, respectively.

**Expression analysis**
Col-0 and *elf3-8* plants were grown under 16 hrs light and 8 hrs dark conditions for 7 days at 22˚C and transferred plants on MS and CIM plates under constant light conditions at 22˚C. Plants were sampled every four hours in duplicate over a twenty-four-hour period resulting in four time courses (Col-0 MS, Col-0 CIM, *elf3-8* MS and *elf3-8* CIM) with twelve samples each. Resulting samples were used for RNA extraction (RNeasy Plant mini kit (Qiagen)), and Illumina RNA sequencing libraries (NEBnext Ultra II RNA Library Prep kit, (New England Biolabs)) were prepared. The libraries were quality controlled and sequenced on an Illumina NextSeq 500 platform (Illumina). The resulting sequence was evaluated for quality and then mapped to the TAIR10 reference genome (www.arabidopsis.org/) with transcripts per million (TPM) used for downstream analysis. The samples were analyzed for cycling behavior as previously described

(*21*); briefly, the time courses were formatted for analysis with the super cycling pipeline (https://gitlab.com/NolanHartwick/super_cycling) (*20*), which leverages a model-based approach to identify cycling genes called HAYSTACK (*23*). Cycling parameters such as period, phase and amplitude were estimated and used to identify cycling genes in the dataset (Data S2).
For RT-qPCR analysis, RNA was isolated from whole roots of 7 day-old Col-0, *elf3-8*, and *lnk1*/*lnk2* with 16 hrs light and 8 hrs dark conditions treated with MS and CIM for 0, 8, 16, 24 hrs under continuous light condition using the RNeasy Plant kit (QIAGEN). First-strand cDNA was synthesized using the ReverTra Ace qPCR Master Mix with gDNA Remover (TOYOBO Co., Ltd). RT-qPCR was performed using the THUNDERBIRD SYBR qPCR Mix (TOYOBO) on a real-time PCR Eco system (PCRmax). The primers used in this study are listed in Table S1. The RT-qPCR efficiency and $C_T$ value were determined using the standard curves for each primer set. Efficiency-corrected transcript values of three biological replicates for all samples were used to determine relative expression values. Each value was normalized against the level of *IPP2* whose expression is not cycling under continuous light conditions (*45*).

**ChIP-seq assay**
ChIP-seq to identify DNA regions bound by ELF3 complex was performed with several modifications to the enhanced ChIP-seq (eChIP-seq) protocol described in (46, 47). Over 2,000 seedlings per wild-type (Col-0) and ELF3-FLAG over expressor were grown on MS medium overlaid with a 108 µm nylon mesh for 7 days with 16 hrs light and 8 hrs dark conditions. At ZT2 on day 7, the seedlings were transferred to CIM along with the mesh for 24 hrs under continuous light condition. After 24 hrs of callus induction, the aerial parts and roots of the seedlings were dissected, and 400-600 mg of each was collected and immediately frozen in liquid nitrogen. The frozen roots were ground into a fine powder using a mortar and pestle, while keeping them cooled with liquid nitrogen. Collected powders were crosslinked with 1% formaldehyde in phosphate-buffered saline (PBS) buffer containing 1 mM Pefabloc SC (Sigma-aldrich), cOmplete proteinase inhibitor cocktail (Sigma-aldrich), 0.3% Triton X-100, and Ethylene Glycol-bis (Thermo Fisher Scientific) for 10 min in room temperature. A final concentration of 200 mM glycine was applied for 5 min to deactivate the remaining formaldehyde. The fixed tissues were washed with ice-cold PBS buffer twice with centrifugation at 5,000 g for 5 min. The tissue pellet was dissolved by the low-salt ChIP buffer without Triton X-100 [50 mM HEPES-KOH (pH 7.5), 150 mM NaCl, 1 mM EDTA, 0.1% Sodium deoxycholate, and 0.1% SDS] containing cOmplete proteinase inhibitor cocktail to a total volume of 2.2 ml. For sonication, ultrasonic homogenizer VP-050 (TAITEC) was used with 100% power (50 W) and 20 cycles of 15 sec on, 5 sec off in an ice-cold water bath. After sonication, the sample was centrifuged at 20,000 g for 10 min, and Triton-X 100 was added to the collected supernatant to a final concentration of 1%. As a shearing check, a small amount of chromatin (20 µl) was evaluated, and the size range of chromatin was 200-600 bp. The sonicated sample was incubated with a primary antibody (Monoclonal anti-FLAG M2 antibody produced in mouse (Sigma-aldrich), 1:1000) for overnight at 4°C with rotation. The sample containing the antibody-ELF3 complex was incubated with Dynabeads Protein G (Thermo Fisher Scientific) for 2 hrs at 4°C with rotation. For washing, the beads were rotated at 4°C for 10 min in the following buffers: once with low-salt ChIP buffer [50 mM HEPES-KOH (pH 7.5), 150 mM NaCl, 1 mM EDTA, 1% Triton X-100, 0.1% Sodium deoxycholate, and 0.1% SDS] containing cOmplete proteinase inhibitor cocktail, twice with high-salt ChIP buffer [50 mM HEPES-KOH (pH 7.5), 350 mM NaCl, 1 mM EDTA, 1% Triton X-100, 0.1% Sodium deoxycholate, and 0.1% SDS]

containing cOmplete proteinase inhibitor cocktail, once with ChIP buffer [10 mM Tris-HCl (pH 8.0), 250 mM LiCl, 0.5% NP-40, 1 mM EDTA, and 0.1% Sodium deoxycholate], and once with TE buffer. After removing the TE buffer, the ELF3 complex was eluted with 100 µl of ChIP elution buffer [50 mM Tris-HCl (pH 7.5), 10 mM EDTA, and 1% SDS] by incubating at 65°C for 15 min. The eluate was then treated with 4 µl of Proteinase K (20 mg/ml) 55°C overnight. The immunoprecipitated DNA was purified using the Monarch PCR & DNA Cleanup Kit (New England Biolabs). The ChIP-seq libraries were constructed using the ThruPLEX DNA-Seq Kit (Clontech) and purified using SPRIselect Beads (Beckman Coulter). The sequencing was performed by the NovaSeq X Plus sequencer (Illumina). Two independent biological replicates were performed for the ChIP-seq experiments.

Reads were mapped on the *Arabidopsis* TAIR10 genome using Bowtie (*48*). Peak calls were performed using the MACS2 program (*49*) and the "intersect" function of the BEDTools (*50*) was used to determine reproducibly bound regions in the biological replicates. The "closest" function of the BEDTools was used to assign 5 nearest genes per each peak, and among those genes, genes with corresponding peak in -3,000 bp to +1,500 bp from TSS were retained as candidate targets. The consensus sequences of the ELF3-bound regions were identified using MEME-ChIP program (*51*).

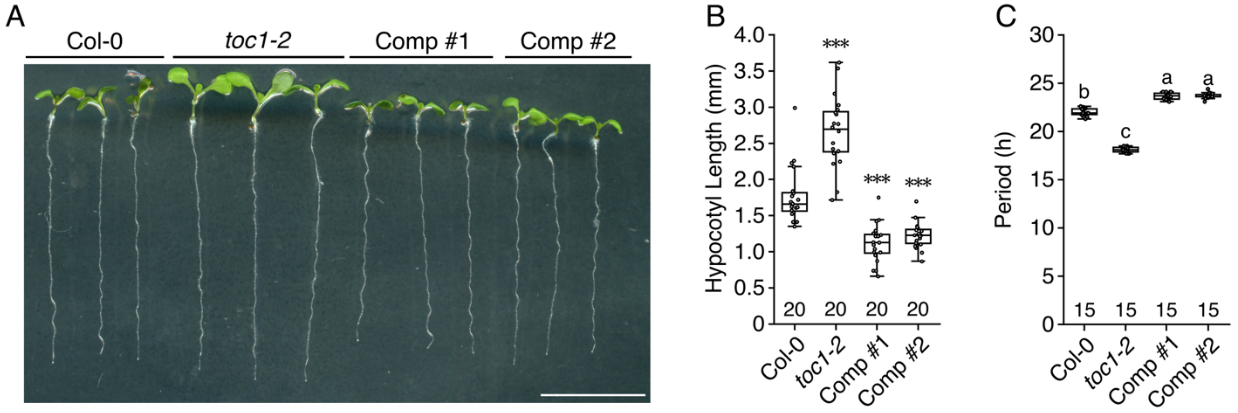


**Fig. S1.**

***pTOC1::TOC1-GFP* construct is functional in the *toc1-2*.** (A) 7-day-old seedlings of *toc1-2* mutant and the complemented line expressing *pTOC1::TOC1-GFP* grown on MS media under LD condition. Scale bars = 13 mm. (B) Hypocotyl length of the plants shown in (A) (n = 20). Significant differences from Col-0 were determined using the Student's *t*-test (****p* < 0.001). (C) LUC intensity of the *pCCA1::LUC* reporter in *toc1-2* and *TOC1-GFP* complemented lines. Seedlings were grown on MS medium under LD condition for 6 days. LUC luminescence intensity was quantified under continuous light condition for 120 hrs. The graph shows the average period of *pCCA1::LUC* intensity from 15 samples. Statistically significant differences between samples determined using the Tukey's honestly significant difference test (*p* < 0.05).

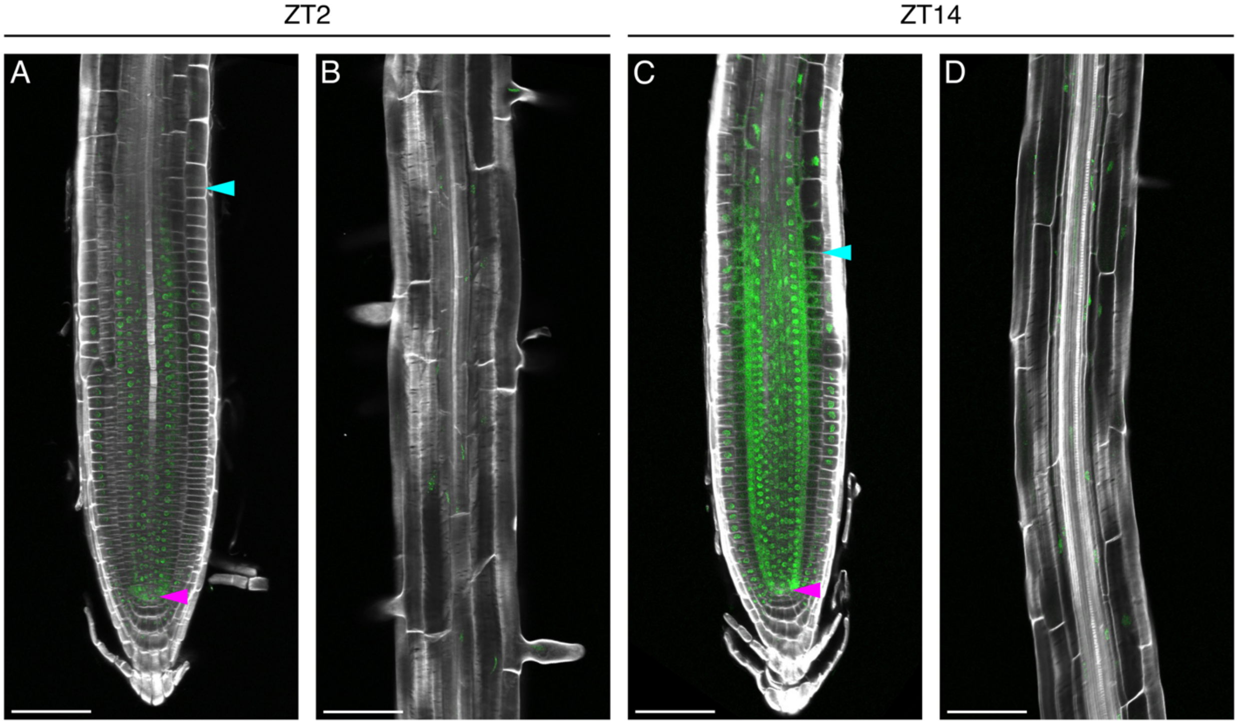


**Fig. S2.**

**TOC1-GFP in primary roots.** *pTOC1::TOC1-GFP*/*toc1-2* at Zeitgeber time (ZT) 2 in the meristem zone (A) and the elongation and maturation zones (B), and at ZT10 in the meristem zone (C) and the elongation and maturation zones (D) of 7-day-old seedlings grown on MS medium. Roots were stained with SR 2200 dye. Pink arrow heads indicate the position of the quiescent center. Blue arrowheads indicate the end of the meristem zone. Scale bars = 75 µm. GFP signals are displayed with a linear adjustment (minimum 0; maximum 35,000) applied uniformly for visualization only.

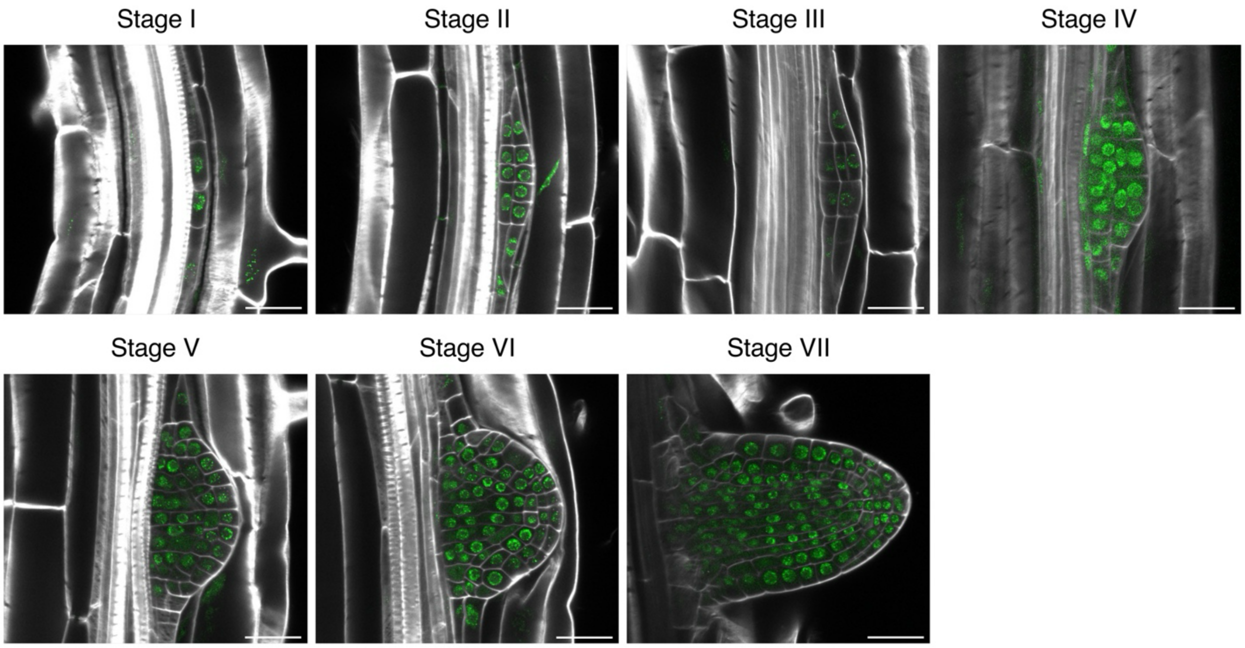


**Fig. S3.**

**TOC1-GFP in lateral root primordia (LRP).** TOC1-GFP expression at ZT10 in each stage of LRP in *pTOC1::TOC1-GFP*/*toc1-2* seedlings grown on MS medium for 10 days. Roots were stained with SR2200 dye. Scale bars = 25 µm. GFP channel images are shown using a uniform linear display range (minimum 0; maximum 30,000) for visualization purposes.

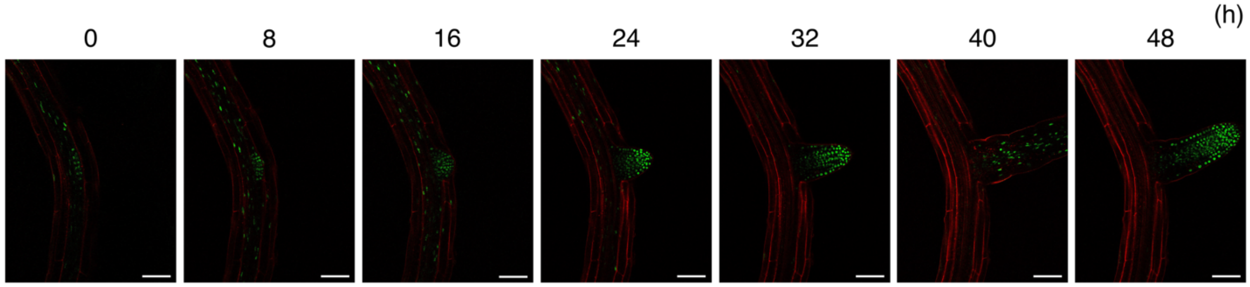


**Fig. S4.**

**The TOC1-GFP fluorescence in the cells of maturation zone and LRP from the 7-day-old seedlings grown on MS medium.** *pTOC1::TOC1-GFP*/*toc1-2* roots were stained with PI solution. Images were taken every 8 hrs for 48 hrs. Scale bars = 100 µm. GFP images are displayed with identical linear brightness settings (minimum 5,000; maximum 30,000) applied uniformly across panels.

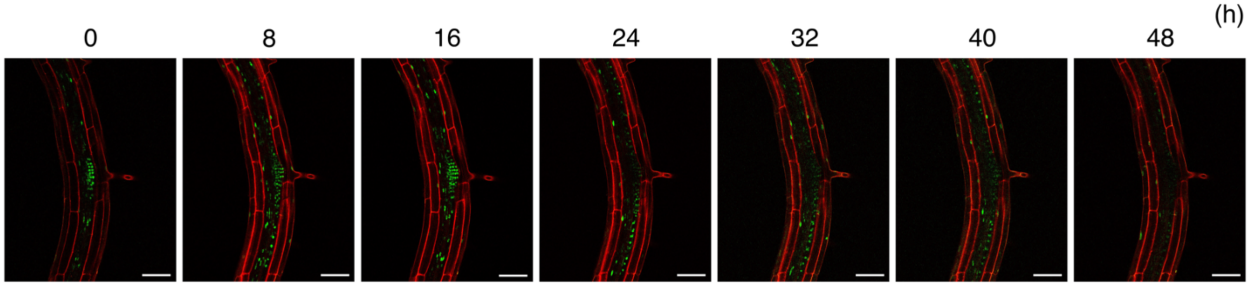


**Fig. S5.**

**The TOC1-GFP fluorescence in the cells of maturation zone and LRP from the 7-day-old seedlings grown on MS medium transferred to CIM.** *pTOC1::TOC1-GFP*/*toc1-2* roots were stained with PI solution. Images were taken every 8 hrs for 48 hrs.Scale bars = 100 µm. GFP signals are shown using uniform linear display settings (minimum 5,000; maximum 30,000) for visualization only.

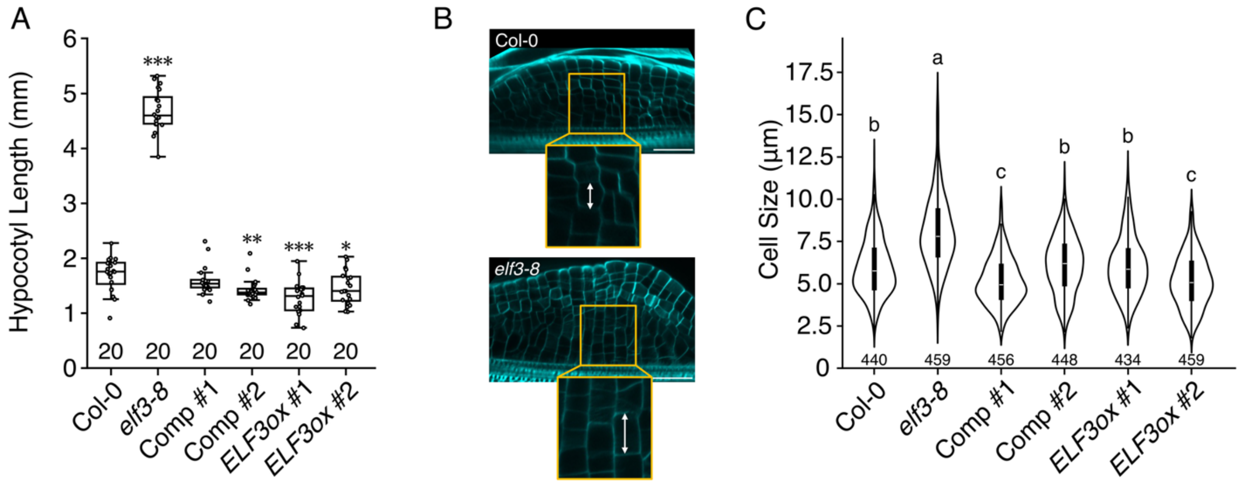


**Fig. S6.**

**Cell size of *elf3-8* was longer than that of Col-0.** (A) Hypocotyl length of Col-0, *elf3-8*, two independent *ELF3* complementation lines, and two independent *ELF3ox* lines (n = 20). Significant differences from Col-0 were determined using the Student's *t*-test (***$p < 0.001$, **$p < 0.01$, *$p < 0.05$). (B) Confocal microscope images of Col-0 and *elf3-8* used for measuring callus cell length. The callus was cleared using ClearSee and stained with Fluorescent Brightener to visualize cell walls. The upper image shows the overall view of the callus (scale bar, 75 µm), and the double-headed arrow in the lower image indicate the measurement of cell length. (C) Cell length in the callus of Col-0, *elf3-8*, two independent *ELF3ox* lines, and two independent *ELF3* complemented lines (n > 400). The number of measured cells is indicated below each element. Statistically significant differences between samples determined using the Tukey's honestly significant difference test ($p < 0.05$).

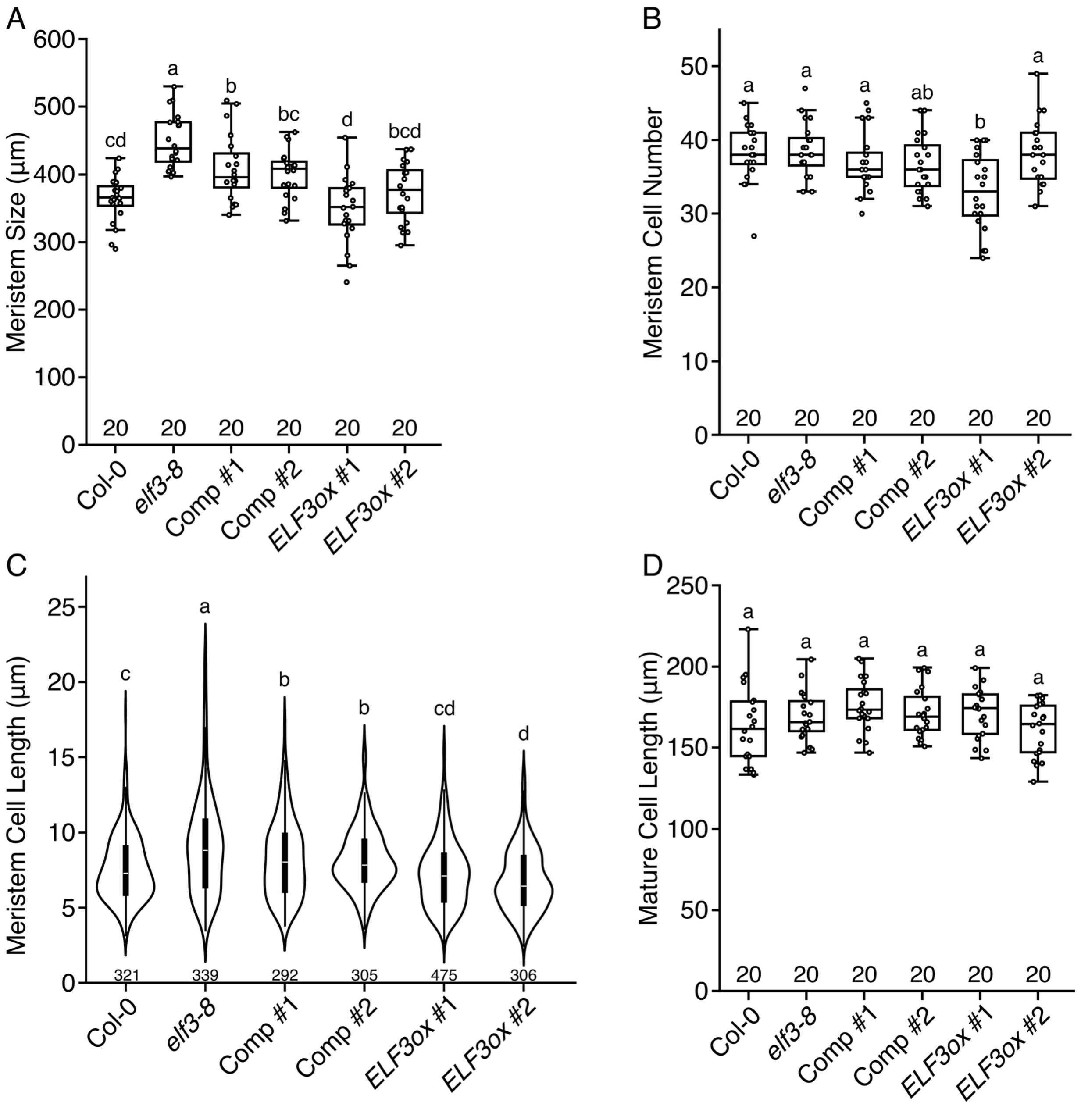


**Fig. S7.**

***elf3-8* exhibits longer root cells in the meristem zone compared to Col-0.** (A) Length of the meristem zone of 7-day-old roots of Col-0, *elf3-8*, two independent *ELF3* complemented lines, and two independent *ELF3ox* lines (n = 20). Statistically significant differences between samples determined using the Tukey's honestly significant difference test ($p$ < 0.05). (B) Cortex cell numbers in the meristem zone of 7-day-old roots of Col-0, *elf3-8*, two independent *ELF3* complemented lines, and two independent *ELF3ox* lines (n = 20). Statistically significant differences between samples determined using the Tukey's honestly significant difference test ($p$ < 0.05). (C) Cortex cell length in the meristem zone of 7-day-old Col-0, *elf3-8*, two independent *ELF3* complemented lines and two independent *ELF3ox* lines (sample sizes are shown below each box). Statistically significant differences between samples determined using the Tukey's honestly significant difference test ($p < 0.05$). (D) Length of mature cortex cells of 7-day-old Col-0, *elf3-8*, two independent *ELF3* complemented lines and two independent *ELF3ox* lines (n = 20). Statistically significant differences between samples determined using the Tukey's honestly significant difference test ($p < 0.05$).

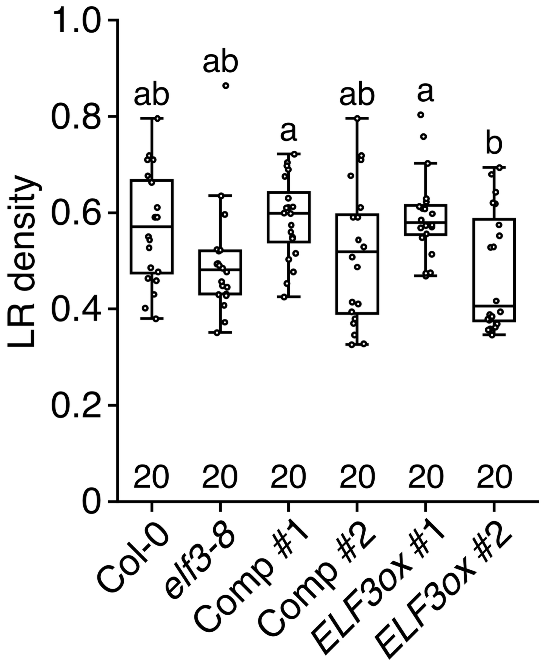


**Fig. S8.**
**No significant difference in whole root LRP density was observed between *elf3-8* and Col-0.** Number of emerged LRs and LRPs per root length in 7-day-old seedlings of Col-0, *elf3-8*, two independent *ELF3* complemented lines and two independent *ELF3ox* lines (n = 20).

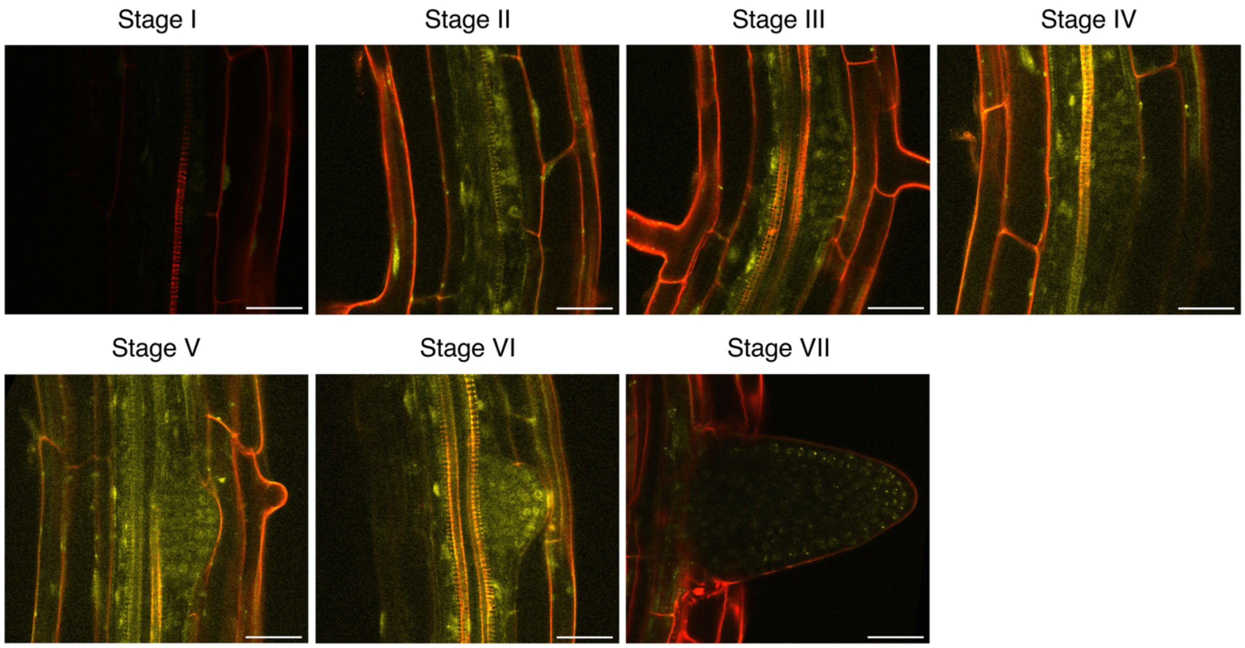


**Fig. S9.**

**YFP-ELF3 expression at ZT12 in each stage of LRP in *pELF3::YFP-ELF3* seedlings grown on MS medium for 10 days.** Roots were stained with PI solution. Scale bars = 25 µm. YFP channel images are displayed with a uniform linear adjustment (minimum 0; maximum 35,000) applied solely to enhance visualization.

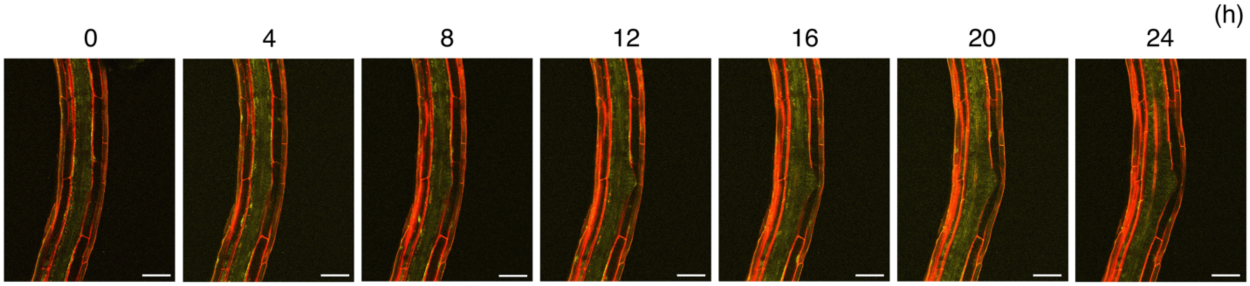


**Fig. S10.**

**YFP-ELF3 expression pattern in CIM treated roots.** Temporal expression pattern of *pELF3::YFP-ELF3* around LRP at 0, 4, 8,12, 16, 20, and 24 hrs after transferring 7-day-old seedlings to CIM. Roots were stained with PI solution. Scale bars, 75 µm. YFP and PI channels are shown using linear display settings (YFP: minimum 0, maximum 35,000; PI: minimum 0, maximum 10,000) applied uniformly for visualization.

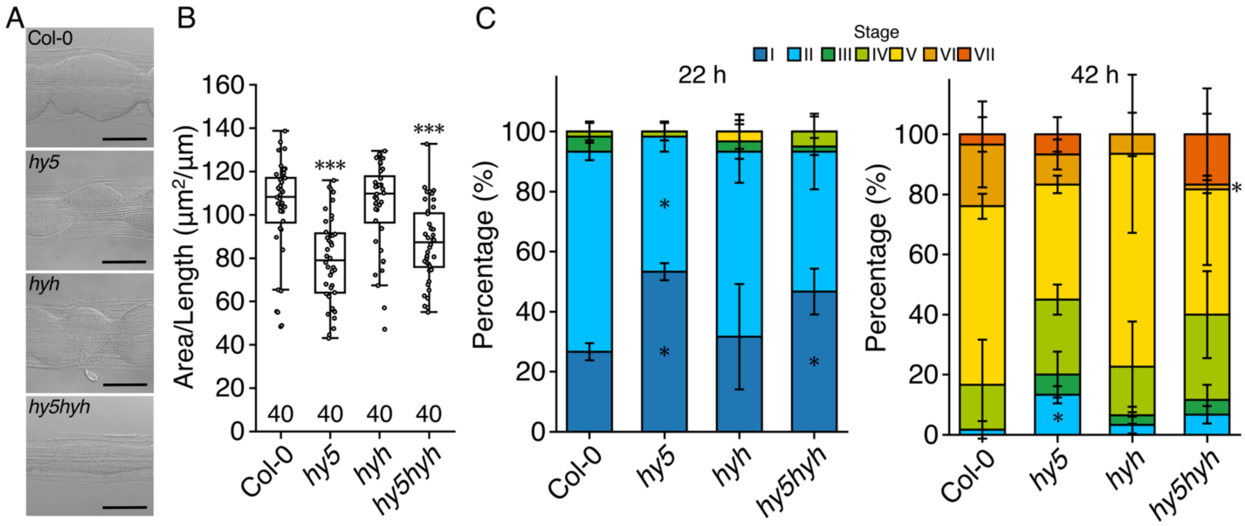


**Fig. S11.**

**HY5 involved in callus and LR development.** (A) Callus images of Col-0, *hy5*, *hyh*, and *hy5*/*hyh* grown under continuous light for three days after being transferred to CIM following 7 days of growth on MS media under LD conditions. Scale bars, 100 µm. (B) Callus area of Col-0, *hy5*, *hyh*, and *hy5/hyh* double mutant grown under continuous light for three days after being transferred to CIM following 7 days of growth on MS media under LD conditions. $^{**}p < 0.01$ determined using Student's *t*-test compared to Col-0 data. (C) LR number in *hy5*, *hyh*, and *hy5*/*hyh* double mutants. Percentage of the LRP at different developmental stages were counted after 22- and 42-hrs of gravistimulation. Data represent mean ± SE from three independent biological replicates, each consisting of 20 seedlings. Statistical significance relative to Col-0 at each stage was determined using a generalized mixed model followed by Holm's P-value adjustment in each stage ($^{**}p < 0.01$, $^{*}p < 0.05$). Colors on the top denote the different LRP stages.

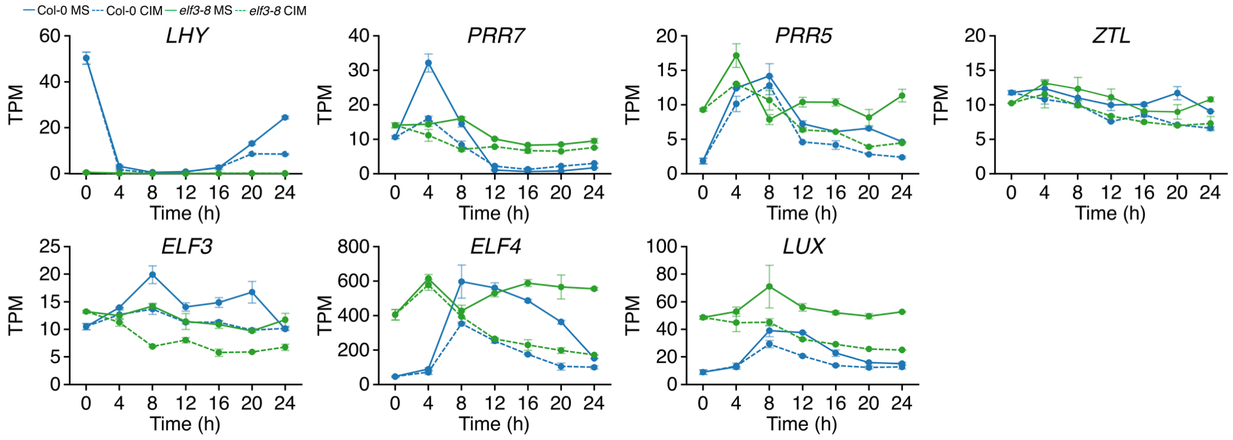


**Fig. S12.**

**Expression patterns of several core-circadian clock genes from the RNA-seq dataset.** All genes expression data are shown with blue representing Col-0 and green representing *elf3-8*; solid lines indicated MS medium, and dashed lines indicate CIM treatment. The X-axis indicates treatment time (hrs), and the y-axis represents expression values with transcripts per million (TPM).

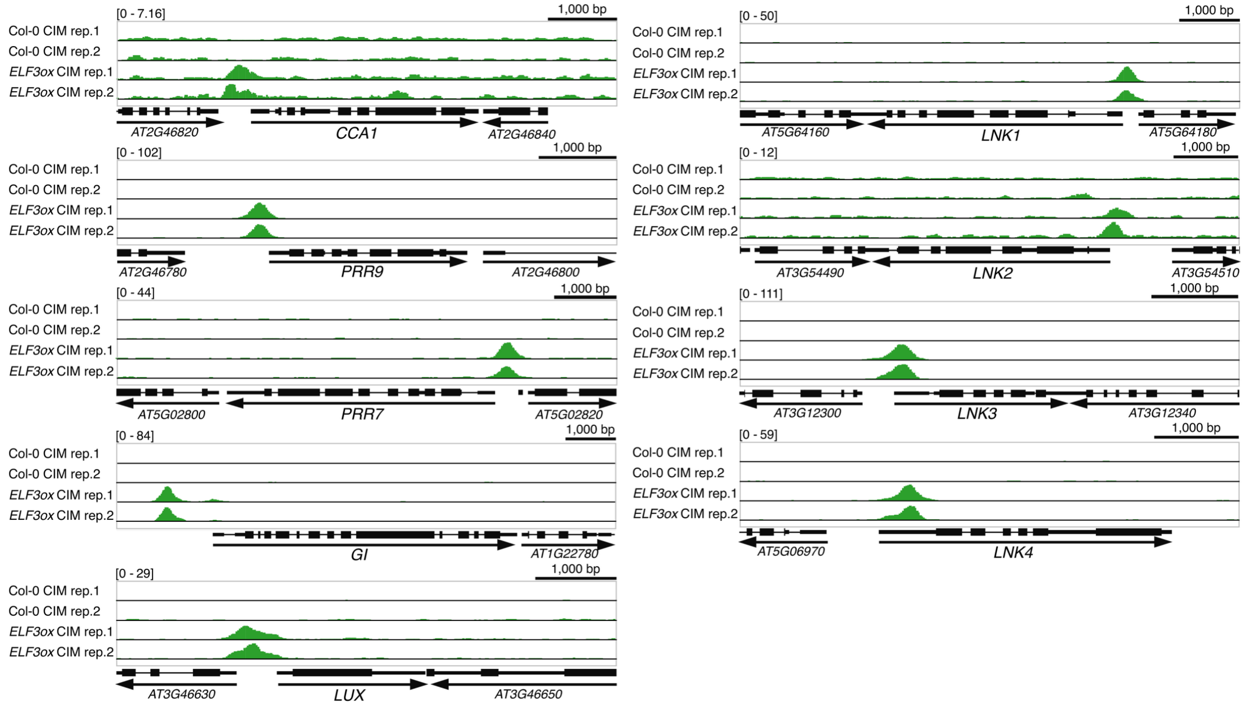


**Fig. S13.**

**Binding of ELF3 to the promoter regions of ELF3 target circadian clock genes.** Visualization of ELF3-FLAG ChIP-seq data using GVI. The promoter region of selected circadian clock genes. "Black boxes" indicate ORFs (thick boxes: exon; lines: intron) and black arrows beneath each ORF represents the direction of transcription. The scale for peak detection on the Y-axis is shown to the left of each graph. The scale bars above graphs indicate 1,000 bp.

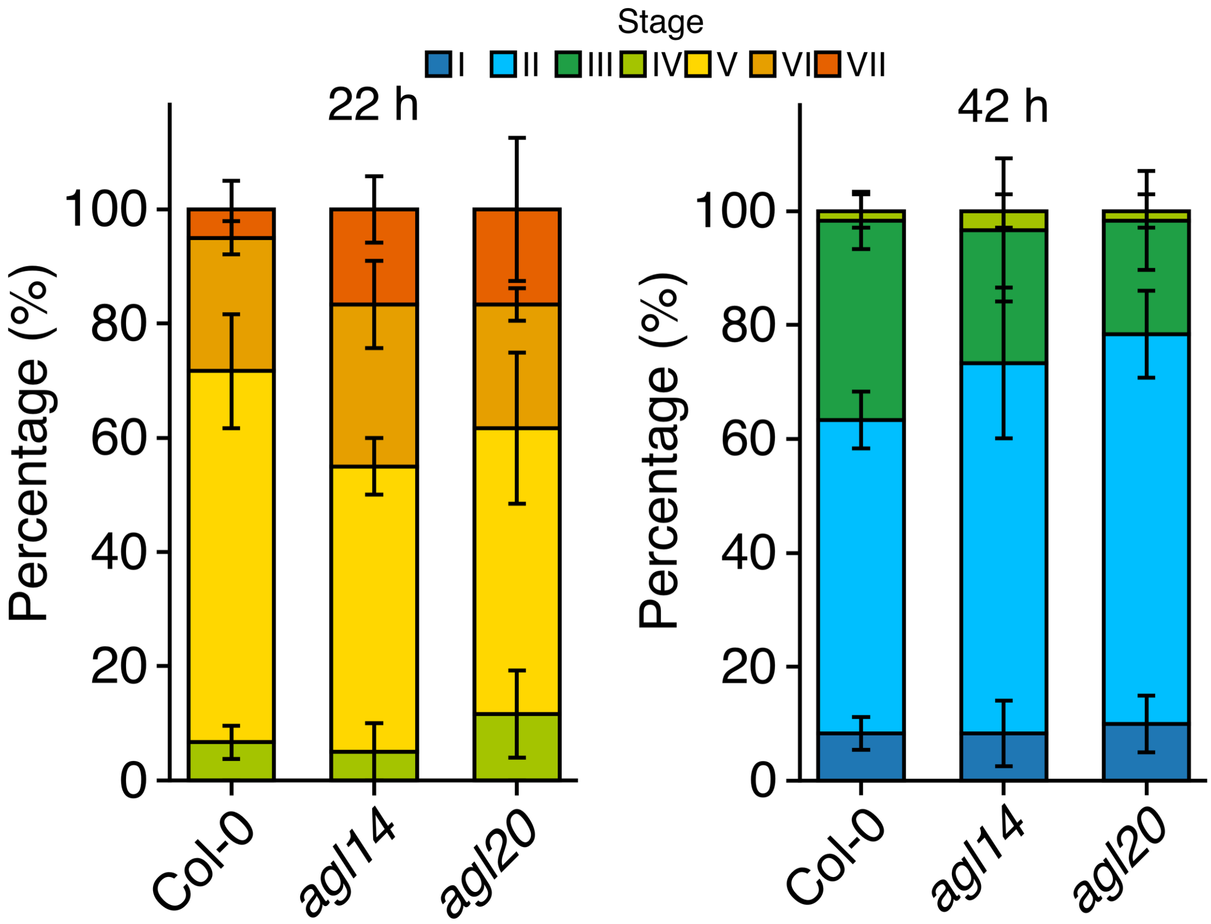


**Fig. S14.**

**LR number in *agl* mutants.** Percentage of the LRP at different developmental stages were counted after 22- and 42-hr of gravistimulation in Col-0, *agl14*, *agl19*, and *agl20*. Data represent mean ± SE from three independent biological replicates, each consisting of 20 seedlings. Colors on the top denote the different LRP stages.

**Table S1.**
**List of primers used in this study.**

| Primer Name | Sequence (5' to 3') | Purpose |
|---|---|---|
| SALK LB | TTTCGCCTGCTGGGGCAAACCAG | Genotyping, SALK T-DNA |
| GABI LB | ATAATAACGCTGCGGACATCTACATTTT | Genotyping, GABI kat T-DNA |
| *agl14* LB | TTTTTAACCCTCTTGGCTTGC | Genotyping, SALK T-DNA |
| *agl14* RB | TTCGACACACACATGTAGCAC | Genotyping, SALK T-DNA |
| *agl20* LB | AAGGATGAGGTTTCAAGCGTC | Genotyping, SALK T-DNA |
| *agl20* RB | TGGCGAATTCATAAAGTTTGC | Genotyping, SALK T-DNA |
| *lnk1* LB | TGTTCCCGTATCACGAGAAAC | Genotyping, SALK T-DNA |
| *lnk1* RB | AGCGGTAGCTGAGGATTTCTC | Genotyping, SALK T-DNA |
| *lnk2* LB | TCCAGTGTTCCATCCATTTTC | Genotyping, GABI kat T-DNA |
| *lnk2* RB | ATTGTTCTATGCGATTGGCAG | Genotyping, GABI kat T-DNA |
| *elf3-8* dCAPs Fw | TCAACTATTGAGTTGCACAGACTGATTAAGG | Genotyping, elf3-8 mutant |
| *elf3-8* dCAPs Rv | TCCGGTGATGCAGCAATAAGTTTTTGAAT | Genotyping, elf3-8 mutant |
| *toc1-2* dCAPS Fw | TTTTTACTCTCCTTTCAGAGTGTTCTTACCAA | Genotyping, toc1-2 mutant |
| *toc1-2* dCAPS Rv | GACAAGAACCTATAAGAGATCTGCAATCTAAAGT | Genotyping, toc1-2 mutant |
| *toc1-2* dCAPS 3'UTR Rv | CAACGGTTTTCAGTTCTTGGTGTATCT | Genotyping, toc1-2 mutant |
| *YFP-cELF3*-Fw | GCTGTACAAGAGCGCTATGAAGAGAGGGAA | ELF3 cDNA region cloning |
| *YFP-cELF3*-Rv | GTGGATCCGGAGCGCTTAAGGCTTAGAGGA | ELF3 cDNA region cloning |
| BamHI-pDonr201-F | GCGCTCCGGATCCACCCAGCTTTCTTGTACAAAG | ELF3 cDNA region for fusing with YFP |
| *CFP*-R | AGCGCTCTTGTACAGCTCGTCCATG | ELF3 cDNA region for fusing with YFP |
| *pELF3*-TA-Fw | AGTATTTTGAACTCCGAAATGATTT | pELF3 cloning |
| *pELF3*-TA-Rv | CACTCACAATTCACAACCTTTTTCA | pELF3 cloning |
| *cAGL14* Fw | TGTACAAAAAAGCAGGCTTTATGGTGAGGGG AAAGACAGAGATG | cAGL14 cloning |
| *cAGL14* Rv | TCCTCGCCCTTGCTCACCATGTTTGAAGGAGG AAACTTTTTGAAGTGTCG | cAGL14 cloning |
| *cAGL20* Fw | TGTACAAAAAAGCAGGCTTTATGGTGAGGGGC AAAACTCAGA | cAGL20 cloning |
| *cAGL20* Rv | TCCTCGCCCTTGCTCACCATCTTTCTTGAAGAAC AAGGTAACCCAATGAAC | cAGL20 cloning |
| *LUX*crispr-Fw | TTGGGTCTCAATTGATTCCACCGAATTTGGCGAG TTTTAGAGCTAGAAATAGCA | crispr guideRNA for lux |

| | | |
|---|---|---|
| *LUX*crispr-Rv | TTGGGTCTCTAAACGAATCGTACGGCTTCTCTCC TGCACCAGCCGGGAATCGAA | crispr guideRNA for lux |
| *pLNK1* Fw | CACCGATTCTTTGGGCTTTCTTCGGC | pLNK1 cloning |
| *pLNK1* Rv | ATTGTTGTCACTTGTTACAACTTCTGC | pLNK1 cloning |
| *pTOC1* Fw | GCAGGCTCCGCGGCCGCCGAGATCGCTCGGCTC AACAACAATATATATTC | pTOC1 cloning |
| *pTOC1* Rv | AGCTGGGTCGGCGCGCCAAGTTCCCAAAGCATC ATCCTGAGGAG | pTOC1 cloning |
| *cELF3-FLAG* for pBS Fw | CAGGGCCCCCCCTCGAGGTCGACTATGAAGAGA GGGAAAGATGAGGAGAAGAT | ELF3 cDNA region cloning |
| *cELF3-FLAG* for pBS Rv | ACCGTCATGGTCTTTGTAGTCCATGGTAGGCTTA GAGGAGTCATAGCGTTTACG | ELF3 cDNA region cloning |
| *cELF3-FLAG* for pSK Fw | TGGAGAGAACACGGGGGACTCTAGAATGAAGAG AGGGAAAGATGAGGAGAAGATATTGGAAC | ELF3 cDNA region cloning |
| *cELF3-FLAG* for pSK Rv | CGATCGGGGAAATTCGAGCTGCGGCCGCTTTACT TGTCGTCATCGTCTTTGTAGTCGATGTCAT | ELF3 cDNA region cloning |
| *IPP2* Fw | GAGACGTCTCATGTTTGAGGATG | qPCR |
| *IPP2* Rv | GGAGGAGCAACTCATACTTCGAG | qPCR |
| q*AGL14* Fw | ACTTACTGAATTTAGCAGTTGCAT | qPCR |
| q*AGL14* Rv | TGCAAAGTTGGGTGAGACGA | qPCR |
| q*AGL20* Fw | TAGCAGTACTGAGAGTGTATAAGGA | qPCR |
| q*AGL20* Rv | AGATACATAACACAAGCAGTTCAGA | qPCR |

**Movie S1.**

**Time-lapse imaging of *pTOC1::TOC1-GFP/toc1-2* treated with MS control medium.** 7 days-old seedlings of *pTOC1::TOC1-GFP/toc1-2* under 16 hrs light/8 hrs dark conditions were transferred from the MS medium to chambered cover glass in MS media under continuous light conditions. Time-lapse images were captures using LAS X every 20 mins for 48 hrs. The timestamp in the upper right represents time zero, corresponding to the start of imaging.

**Movie S2.**

**Time-lapse imaging of *pTOC1::TOC1-GFP/toc1-2* treated with CIM.** 7 days-old seedlings of *pTOC1::TOC1-GFP/toc1-2* under 16 hrs light/8 hrs dark conditions were transferred from the MS medium to chambered cover glass in CIM under continuous light conditions. Time-lapse images were captures using LAS X every 20 mins for 48 hrs. The timestamp in the upper right represents time zero, corresponding to the start of imaging.

**Movie S3.**

**Time-lapse imaging of *pELF3::YFP-ELF3/elf3-8* treated with CIM.** 7 days-old seedlings of *pELF3::YFP-ELF3/elf3-8* under 16 hrs light/8 hrs dark conditions were transferred from the MS medium to chambered cover glass in CIM under continuous light conditions. Time-lapse images were captures using LAS X every 20 mins for 48 hrs. The timestamp in the upper right represents time zero, corresponding to the start of imaging.

**Data S1. (separate file)**

Expression and cycling calls for the Col-0 and *elf3-8* root time courses on CIM and control media (MS).

**Data S2. (separate file)**

Gene Ontology (GO) overrepresented terms for genes cycling in Col MS and *elf3-8* CIM, but not under Col CIM or *elf3-8*.

**Data S3. (separate file)**

Expression patterns and cycling calls for 91 clock, flowering and light signaling genes.

**Data S4. (separate file)**

*ELF3*-associated peaks located within 3,000 bp upstream of gene coding regions.

**Data S5. (separate file)**

Genes that were responsive to CIM treatment independently of the ELF3 genetic background.
